# Dynamics of an Expanding Black Rhinoceros (*Diceros Bicornis Minor*) Population

**DOI:** 10.1101/018523

**Authors:** Peter R Law, Brad Fike, Peter C. Lent

**Affiliations:** Centre for African Conservation Ecology, Nelson Mandela Metropolitan University, P.O. Box 77000, Port Elizabeth, 6031, Republic of South Africa; P.O. Box 4038, Rosehill Mall, Port Alfred, 6170, Republic of South Africa; Department of Zoology and Entomology, University of Fort Hare, Alice, 5700, Republic of South Africa

**Keywords:** black rhinoceros, density dependence, matrix models, megaherbivores, population dynamics, transient dynamics, stochasticity

## Abstract

Population dynamics is a central component of demography and critical for meta-population management, especially of endangered species. We employed complete individual life records to construct census data for a reintroduced black rhinoceros population over 22 years from its founding and investigated that population’s dynamics to inform black rhinoceros meta-population management practice and, more generally, megaherbivore ecology. Akaike’s information criterion applied to scalar models of population growth based on the generalized logistic unambiguously selected an exponential growth model (*r* = 0.102 ± 0.017), indicating a highly successful reintroduction, but yielding no evidence of density dependence. This result is consistent with, but does not confirm, the threshold model of density dependence that has influenced black rhinoceros meta-population management. Our analysis did support previous work contending that the generalized logistic is unreliable when fit to data that do not sample the entire range of possible population sizes. A stage-based matrix model of the exponential population dynamics exhibited mild transient behaviour. We found no evidence of environmental stochasticity, consistent with our previous studies of this population that found no influence of rainfall on demographic parameters. Process noise derived from demographic stochasticity, principally reflected in annual sex-specific recruitment numbers that differed from deterministic predictions of the matrix model. Demographically driven process noise should be assumed to be a component of megaherbivore population dynamics, as these populations are typically relatively small, and should be considered in managed removals and introductions. We suggest that an extended period of exponential growth is common for megaherbivore populations growing from small size and that an increase in age at first reproduction with increasing population size, manifest in the study population, may provide a warning of density feedback prior to detectable slowing of population growth rate for megaherbivores.

## INTRODUCTION

Reintroduction is an important strategy of conservation science (Seddon *et al*. 2007) and a critical component of black rhinoceros (*Diceros bicornis*) meta-population management (Emslie 2001). Since reintroductions are sourced from existing populations, understanding population dynamics is vital to conservation theory and practice as regards both the expected performance of the introduced population and possible effects on the harvested population (Armstrong and Seddon 2007). Similarly, understanding the dynamics of populations affected by poaching is important (Brodie *et al*. 2011).

Scalar models of density dependence are prominent in population studies of large herbivores (Owen-Smith 2010). Verhulst (1838) considered the equation *N*′ = *rN* − *ϕ*(*N*) and in particular the form *ϕ*(*N*) = *bN* ^*θ*^ (though with different notation), in the context of population growth. This latter has also been employed as a growth equation for organisms. In that context Nelder (1961) cited Pütter (1920) and Richards (1959) and noted that the equation can be derived from von Bertalanffy’s growth equation (e.g., Bertalanffy 1957). Gilpin and Ayala (1973) generalized the Lotka-Volterra predator-prey equations by replacing *N*/*K* by (*N*/*K*)^*θ*^ in the logistic equation. Gilpin *et al*. (1976) attributed this generalized logistic to Verhulst (1838). Recognition that the linear decline in per capita growth rate (pgr) of the logistic model is unrealistic for populations of large vertebrates (Fowler 1981, 1987; McCullough 1992) led to favouring of the generalized logistic

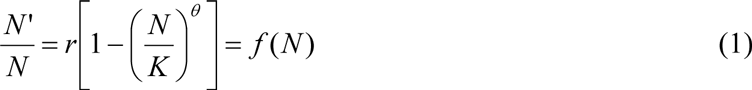

as a flexible model of density dependence, the parameter *θ* controlling the approach to equilibrium *K*, with a value larger than one considered appropriate for large vertebrates, and *r* the intrinsic rate of growth. Fowler (1981) actually only wrote down the logistic rather than the generalized logistic but referred to generalized growth models and cited Richards (1959), Pella and Tomlinson (1969), and Gilpin *et al*. (1976).

The logistic equation (*θ* = 1) can be solved by direct integration using the method of partial fractions. Rewrite the ODE (ordinary differential equation) as

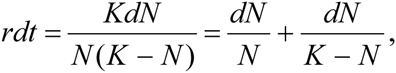

and integrate to obtain, after some manipulation,

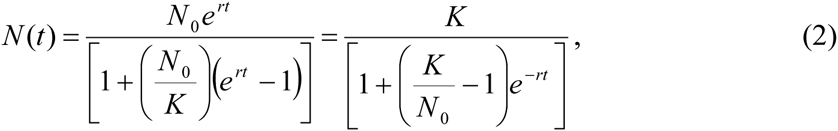

where *N*_0_ = *N*(0). The generalized logistic can be solved in the same manner, or one can put *M* = *N^θ^*, *J* = *K^θ^*, and *s* = *rθ* and the generalized logistic ODE becomes

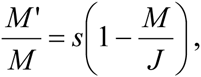

i.e., the logistic equation for these quantities. Hence, the solution of the generalized logistic can be obtained directly from (1) as

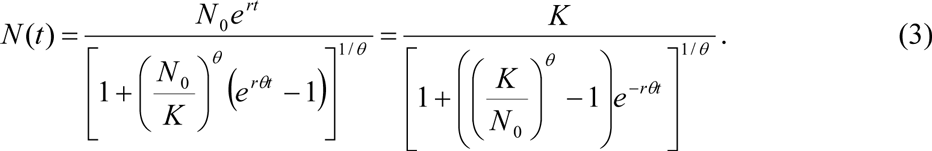

Standard theory of first –order ODEs (existence of a phase flow) guarantees that if one writes the solution (2) in the form *N* (*t*) = *F*(*t*, *N* (0)), then

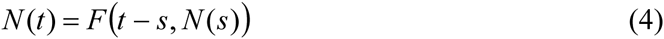

for any *s*, 0 ≤ *s* < *t*. Equation (3) is easily confirmed for the generalized logistic by algebraic substitution in (2).

From (2), one sees that for *N*_0_ < < *K* and so that (*N*_0_/*K*) < *e*^*-rt*^, then for large *θ* the denominator in the first form is approximately one and *N*(*t*) behaves as exponential growth over that range of *t* values. Yet the solution still converges on the equilibrium *K* for large *t* and does so rapidly once the rate of growth begins to decline. Indeed, as *θ* → ∞, the solution converges on exponential growth until *N*(*t*) reaches *K* and growth ceases, which is an extreme form of threshold model. McCullough (1999) proposed that pgr for large herbivores might remain constant from low abundance to near equilibrium (corresponding to exponential growth) and then decline rapidly as equilibrium is approached. For such a model, *N*(*t*) is continuous but only piecewise differentiable, with a point of nondifferentiability at the threshold value *N*\* at which exponential growth ceases. If the decline is modelled by the generalized logistic, then the expression for *N*(*t*) is an exponential growth curve joined at *N*\* to a solution of the generalized logistic with the same *r* and suitable *θ* and *K* (see also Owen-Smith 2010:39). Taking *θ* = 1 gives linear decline in pgr after the threshold, sometimes called the ‘ramp’ model.

The graph of (2) is sigmoid with point of inflexion at

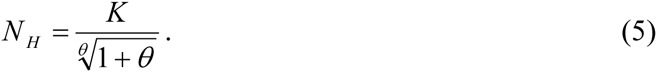

*N*_*H*_ is also the abundance at which population growth rate *N*′ (*t*) is a maximum. For *θ* > 1, *N*_*H*_ > *K*/2 and approaches *K* as *θ* → ∞. Hence, for *θ* > 1, the abundance for optimal sustainable harvesting is nearer to *K* than the value of *K*/2 for the logistic. For a threshold model of population growth, optimal sustainable harvesting can be achieved by harvesting at the model’s threshold value *N*\*. Thus, the appropriate model of population dynamics informs not only how the population approaches equilibrium but also how harvesting can be practised. In particular, Emslie (2001) advocated an approach to meta-population management for black rhinoceros based on a threshold model of density dependence but the scarcity of undisturbed expanding populations poses an obstacle to testing that assumption. One of our objectives was to see whether our study population of black rhinoceros (*Diceros bicornis minor*) conformed to expectations based on a threshold model of dynamics.

Ideally, fitting a time series of abundances to the solution (2) of (1) by nonlinear regression yields estimates of the model parameters by maximum likelihood. Unfortunately, time series of data fit to (2) may not yield precise estimates for *r*, *K*, and *θ* when the data does not adequately cover the full range of values from near zero to *K*; *θ* in particular is often imprecisely estimated (Nelder 1961). Time series of abundances are often modelled with discrete-time versions of (1); when the data fluctuates about a presumed equilibrium but does not sample the population at low abundance, maxima of the likelihood may be determined by the product *rθ* rather than these parameters separately, entailing a redundancy in *r* and *θ* and non-uniqueness of estimates (Polansky *et al*. 2009; Clark *et al*. 2010). Moreover, there are several ways to discretize a continuous-time model and various forms appear in the literature. Furthermore, a discrete-time model may misrepresent continuous-time dynamics, or discrete-time dynamics may be misrepresented by an inappropriate time step in the model, in each case yielding erroneous estimates of *θ* (Doncaster 2008). Scalar models themselves misrepresent population dynamics by ignoring population structure, which may obscure transient dynamics (Koons *et al*. 2005). Complicating matters further are the influences of environmental stochasticity and, for small populations such as reintroduced populations, demographic stochasticity. Hence, extracting useful information from a time series of abundances faces a variety of challenges.

Our dataset consisted of a time series of censuses of a black rhinoceros population that grew monotonically from its reintroduction in 1986 through the end of 2008 without reaching equilibrium. We fitted scalar models of population growth to these data to evaluate whether density dependence acted during this time and if so in what form. This exercise addressed the objective stated above regarding the probing of population dynamics of black rhinoceros in particular, and megaherbivores in general, but also permitted us to explore the difficulties mentioned in the preceding paragraph of prescribing and fitting models to abundance data. Our data exemplified the opposite density extreme to that considered by Polansky *et al*. (2009) and Clark *et al*. (2010), i.e., growth from low numbers rather than populations near equilibrium.

On the basis of the results of the scalar population modelling, we next built a matrix model to assess the relevance of population structure to the dynamics and, in particular, to detect transient dynamics. We also evaluated the importance of environmental and demographic contributions to process noise (Lande *et al*. 2003), during the period after introductions ceased. That environmental variation can influence population dynamics is well known (Owen-Smith 2010 includes reviews for large herbivores). Demographic stochasticity may be important for reintroductions and perhaps for populations of megaherbivores in general. Few estimates of demographic stochasticity of populations of large mammals appear in the literature.

Our study population has been the subject of various studies focusing on other aspects of black rhino, and megaherbivore, ecology, reflecting not only the importance of the critically endangered black rhinoceros but also the fact that the study population has been undisturbed by poaching, been subject to little management intervention, but monitored at the individual level over the 22 years of the study period, resulting in a rare opportunity to study a natural megaherbivore population. These other studies (Lent and Fike 2003; Ganqa *et al*. 2005; Ganqa and Scogings 2007; van Lieverloo *et al*. 2009; Fike 2011; Law *et al*. 2013, 2014) illuminate the results of this paper. Our paper therefore contributes to the valuable study of this particular population and contributes to the understanding of megaherbivore population dynamics (Cromsigt *et al*. 2002; Gough and Kerley 2006; Chamaillé-Jammes *et al*. 2008; Okita-Ouma *et al*. 2010; Owen-Smith 2010; Brodie *et al*. 2011) both for theoretical ecology and conservation science.

## STUDY POPULATION AND DATASET

We base black rhinoceros demography on biological states rather than age (Law and Linklater 2014) and employ the definitions of ‘calf’, subadult’, ‘female adult’ and ‘male adult’ of Law *et al*. (2013, 2014) and Law and Linklater (2014), summarized in Table 1.

**Table 1.** Definitions of biological life stages for demography of the black rhinoceros

| Life Stage | Definition |
| --- | --- |
| Calf | From birth to: observed separation from the mother; at the birth of the mother’s next calf; or the calf’s 4<sup>th</sup> birthday; whichever comes first |
| Subadult | From ceasing to be a calf until becoming an adult |
| Female Adult | The subadult-adult transition occurs for females at first calving or at the 7<sup>th</sup> birthday, whichever comes first |
| Male Adult | The subadult-adult transition occurs at the 8<sup>th</sup> birthday. |

The Great Fish River Nature Reserve (GFRNR), Eastern Cape Province, South Africa, is split into halves by the Great Fish and Kat rivers, and is considered excellent black rhinoceros habitat (Ganqa *et al*. 2005; Ganqa and Scogings 2007; van Lieverloo *et al*. 2009; Fike 2011). Black rhinoceros (*Diceros bicornis minor*) populations were independently introduced into each half of the reserve. The population in the 220 km^2^ western sector (former Sam Knott and Kudu Reserve) is the older, larger, and more consistently monitored of the two, and has been the focus of considerable study, as noted in the Introduction; we refer to it from its founding in June 1986 through December 2008 as the SKKR population (or simply SKKR).

The SKKR population was founded with the release in June, 1986, of one adult female (aged about 24), one subadult (SA) female (aged about three), and two males, both judged to be adults. One of the males died in 1988 from an injury that occurred prior to importation and was treated as a failed import, i.e., excluded from the study.

In October/November 1989, when population size was four, a second cohort was released consisting of three SA females and three SA males, of which one female and one male died in 1989 and another male died in 1990, each regarded as a failed import. In November, 1990, when population size was eight, two adult females (one aged about 15; the other aged about 7 and pregnant) and a SA male were imported. The pregnant female calved in December 1990 but both mother and calf were dead by the end of 1991 and are treated as failed imports. In January 1992, when population size was 11, a female SA and a male adult were released. Between September 1997, when the population was 26, and December 1997, a cohort of 7 females and 6 males, all SAs and all quite young (about three years of age) except one (aged about six) was imported. See Fike (2011).

In summary, 13 males and 15 females were introduced but 3 males and 2 females died soon after release and did not contribute to the population. The surviving imports included only two females and two males that were already adults. One further individual, a female adult, entered the SKKR population, from the eastern sector of the GFRR, in 2003. This immigrant was the only exception to the demographic isolation of the SKKR population during the study period. She calved for the first time after entering the SKKR population and is included as a member of the SKKR population from her time of entry.

The export of 1 SA male and 4 SA females in May 2006 yielded a sex ratio of 9:10 as a result of imports and exports, after discounting the failed imports. These exports were the first removals from the SKKR population, conducted as part of the meta-population management plan to provide donors for reintroductions elsewhere and maintain high population growth rates by preventing density feedback on population growth rate. More substantial removals were conducted after 2008. Our demographic study of the SKKR population focused on the period from reintroduction through the end of 2008 to obtain the longest study period possible with minimal effects from removals. As detailed below, we accounted for the removals in both scalar and matrix modelling, though in different ways.

The complete absence of poaching in the GFRR, the fact that only five rhinos were removed prior to 2009 by management, and the excellent monitoring of the population at the individual level made this population an excellent opportunity for the study of black rhino ecology in general and the performance of a reintroduced population in particular. See the previously cited literature for further details of the population.

SKKR was monitored under BF’s direction as reserve manager by ground patrols and aerial reconnaissance; each animal was ear notched, and births and deaths recorded as part of this monitoring. No animals were handled for the research reported in this paper. The data collected by BF permitted actual population censuses to be computed for any month from Jun-86 through Dec-08. The only uncertainty in such censuses arises from uncertainties in birth and death dates. At the initial recording of each birth and death, an interval of uncertainty was assigned to reflect the precision of the birth or death date (Fike 2011). The interval of uncertainty, in months, centered on the nominal birth date (*d*), was specified by a value *U* so that the interval of uncertainty was *d* – *U* to *d* + *U*. The values of *U* employed by Fike (2011) were: *U* = 0 (uncertainty in the nominal date at most 1 week); *U* = 1; *U* = 3; *U* = 6; *U* = 12; *U* > 12. For 106 births and 15 deaths, 29 had *U* = 0, 38 had *U* = 1, 32 had *U* = 3, 20 had *U* = 6, 2 had *U* = 12, and none had *U* > 12. We therefore computed censuses semi-annually, every June and December to limit the effects of these uncertainties. We then used the assigned intervals of uncertainties to inspect their impact on the censuses by noting when a rhino was unambiguously present or not and counting the **maximum** possible number of rhino that might be added to or subtracted from the nominal census count due to the uncertainty of birth and death dates. Expressed as percentages of the nominal census counts, only three possible modifications exceeded 10%, each at very low population levels (viz., an ambiguity of one rhino in a count of three or four). From Jun-86 through Jun-98, 17 of 25 censuses possessed no ambiguity at all. From Dec-98 through Dec-08, an ambiguity was always present but in 14 of 21 censuses was less than 5%. It was clear that this data could not be modelled as independently and identically distributed. As these were maximum estimates of uncertainty, and ambiguous presence and ambiguous absence, when both present, would tend to cancel, taking the nominal censuses as actual population censuses seemed plausible.

In particular, we considered our data not to include the kind of observation error that arises from employing sample data or count estimates, as many studies are forced to do. We therefore did not need to be concerned with false signals of density dependence (Shenk *et al*. 1998; Freckleton *et al*. 2006) or with conflating sampling error and process noise (deValpine and Hastings 2002).

SKKR grew monotonically on a semi-annual basis to reach 110 (26 calves, 39 subadults, and 45 adults) at the end of 2008. Our dataset consists of population censuses each June and December, from June, 1986, through December, 2008, Fig. 1, computed from complete population records.

## ANALYSES AND RESULTS

### 1. Scalar Population Models

We began by fitting the census data to scalar models of population growth. Since the data in Fig. 1 does not indicate an equilibrium or obvious threshold, we could not include a threshold model amongst the candidate models, e.g., a model based on the ODE of the form

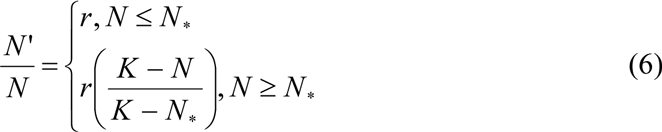

in which it is assumed that *N*_*_ is close to *K*, but threshold-like dynamics can be approximated by large values of *θ* in the generalized logistic.

**Figure 1.**
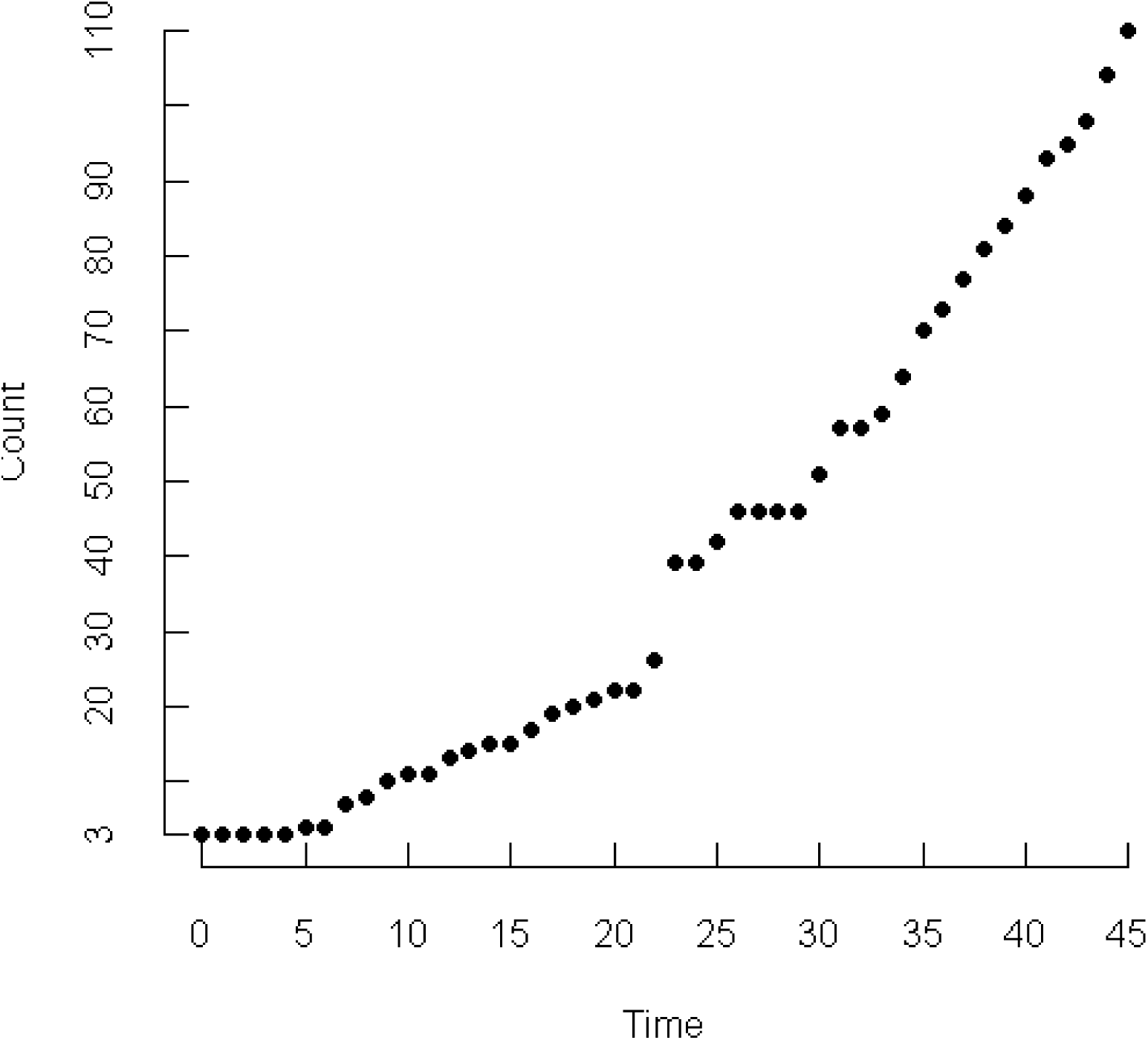
46 **s**emi-annual census counts of the SKKR population from Jun-86 through Dec-08.

As black rhinoceros are aseasonal breeders their population dynamics lack a natural time step, suggesting continuous-time models are appropriate. We write a per capita growth equation like (1) and its solution (2) in the form

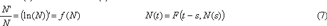

0 ≤ *s* < *t*. Denote the census data at time *t* by *N*_*t*_. Nonlinear regression fits this data to the solution by putting *ε* _*t*_ = *N*_*t*_ − *F* (*t*, *N* _0_), with *ε*_*t*_ independent N(0,*σ*^2^) variates. Since the projected value at *t* depends only on *N*_0_ in this model, *ε*_*t*_ does not participate in the dynamics and so is not a realistic model of process noise. We included this naïve model in our analyses as a way of evaluating the importance of process noise.

In a discrete-time model, as the projected value at the next time step depends on the current value, deviations from the deterministic model participate in the dynamics and can be interpreted as process noise in a straightforward fashion. A semi-annual time step should preclude a pronounced artificial time lag in discrete-time models of population dynamics for black rhinoceros. We considered three discretizations of (7) (Turchin 2003:52). The per capita growth rate 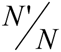 can be discretized as (*N*_*t*+1_ – *N*_*t*_)/*N*_*t*_ to give

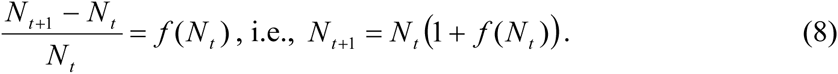

For the generalized logistic, however, this model will produce a negative value for *N*_*t*+1_ if

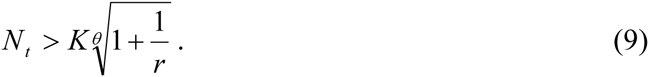

This model is therefore unrealistic for populations for which *r* is large for then *N* may easily exceed this quantity. This flaw may not be an issue for populations of large herbivores, or at least megaherbivores. For *r* = 0.1, say, *N* would have to exceed 11*K* for the logistic to generate negative abundances; for *θ* = 4.5 (a value suggested by Eberhardt *et al*. 2008), *N* would have to exceed 1.7*K*; for *θ* = 10, *N* would have to exceed only 1.27*K*. But even this density may be unlikely in a natural population of large herbivores, so the model may only expose its flaw for large herbivores at rather artificial densities, such as in enclosed populations.

The second discretization notes that 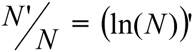 and replaces this quantity by ln(*N*_*t*+1_) – ln(*N*_*t*_) = ln(*N*_*t*+1_/*N*_*t*_) to obtain

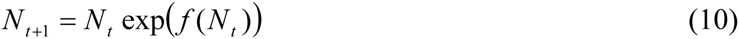

This model cannot generate negative abundances and for that reason is considered more realistic for ecological applications (Turchin 2003:53). Note that (8) can also be obtained from (10) by taking the linear approximation to the exponential function, whence (8) and (10) will be similar when *f*(*N*_*t*_) is small, i.e., when *N*_*t*_ is near *K*.

The third approach to discretization puts

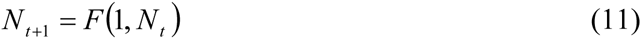

i.e., *N*_*t*+1_ is projected one time step from an abundance of *N*_*t*_ using the solution *F* of the continuous-time model (7). For the generalized logistic, one obtains

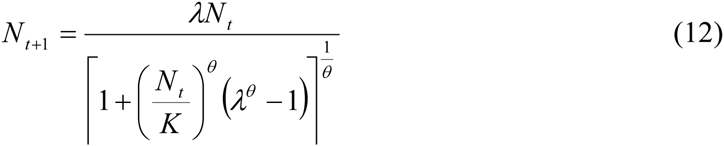

where *λ* = *e*^*r*^. When *θ* = 1, this model is of the form of the Beverton-Holt model; if one expands the denominator using the binomial expansion, the linear approximation is of the form of the generalized Beverton-Holt model

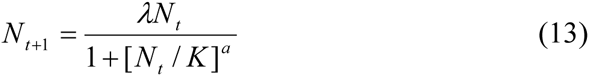

studied by Getz (1996). When *f* is as in the generalized logistic (1), we refer to the model resulting from (8) as the discrete generalized logistic (DGL), the model resulting from (10) as the generalized Ricker (genRicker), and the model (12) resulting from (11) as the stepwise generalized logistic (SGL). The DGL figured prominently in Cromsigt *et al*. (2002) and Okita-Ouma *et al*. (2010). The SGL should be most faithful to continuous-time dynamics.

Continuous-time exponential growth can be obtained from the generalized logistic by setting *K* = ∞, so that *f*(*N*) = *r*. For (8) one obtains *N*_*t*+1_ = *N*_*t*_(1 + *r*), and for (10) and (12) one obtains *N*_*t*+1_ = *N*_*t*_*e*^*r*^, which are just two versions (with different interpretations of *r*) of the same model. We used the latter form.

### Error Structure and Process Noise

When fitting a scalar model of population growth to a time series of census data by nonlinear regression, residuals can be interpreted as process noise in the dynamics. For any deterministic, discrete-time model of the form *N*_*t*+1_ = *G*(*N*_*t*_), if in fact *N*_*t*+1_ = *G*(*N*_*t*_) + *ε*_*t*_ the terms *ε*_*t*_ contribute to the dynamics at each time step in the sense that the projection to time *t*+2 is based on N_*t*+1_, which includes *ε*_*t*_, and thus models process noise as additive on abundance. Unlike these discrete-time models, we noted above that in the continuous-time model *N*(*t*) = *F*(*t*,*N*_0_) + *ε*_*t*_, the term *ε*_*t*_ at time *t* only corrects the projection at time *t* but does not influence the projection at future times and in this sense is not part of the dynamics and so does not realistically model process noise. An error structure additive on abundance, as just described, was employed by Cromsigt *et al*. 2002, and Okita-Ouma *et al*. 2010.

With error structure interpreted as process noise, however, it is also common to model the error structure as multiplicative on abundance, whence additive on *X*_*t*_ := ln(*N*_*t*_), (Polansky *et al*. 2009; Clark *et al*. 2010). For example, in the discrete-time exponential model, if *r* is replaced by *r* + *ε*_*t*_, then one obtains

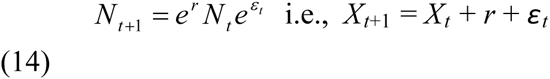

Hence, with process noise additive on vital rates, it is multiplicative on abundance (see Hilborn and Mangel 1997:73–74, Turchin 2003:184). For multiplicative error structure, we therefore wrote *X*_*t*+1_ := ln(*N*_*t*+1_) = ln(*G*(*N*_*t*_)) + *ε_t_*, so if *ε*_*t*_ is N(0,*σ*^2^), exp(*ε*_*t*_) is log-normal.

The models we compared are listed in Table 2.

**Table 2.** Scalar models of population dynamics with two error structures. *N*_*t*_ is the census count at time *t*; *X*_*t*_ = ln(*N*_*t*_); *F*(*t*,N_0_) is the solution of the continuous-time generalized logistic as in (3) and (4); *f* is the per capita growth rate of the generalized logistic as in (1); *ε*_*t*_ are independent N(0,*σ*^2^) variates.

| Model | Additive Error | Multiplicative Error |
| --- | --- | --- |
| Continuous-time<br>exponential (Cexp) | $N_t = N_0 e^{rt} + \varepsilon_t$ | $X_t = X_0 + rt + \varepsilon_t$ |
| Continuous-time<br>generalized logistic (CGL) | $N_t = F(t, N_0) + \varepsilon_t$ | $X_t = \ln(F(t, N_0)) + \varepsilon_t$ |
| Discrete-time exponential (Dexp) | $N_{t+1} = e^r N_t + \varepsilon_t$ | $X_{t+1} = r + X_t + \varepsilon_t$ |
| Discrete generalized<br>logistic (DGL) | $N_{t+1} = N_t (1 + f(N_t)) + \varepsilon_t$ | $X_{t+1} = X_t + \ln(1 + f(N_t)) + \varepsilon_t$ |
| Generalized Ricker<br>(genRicker) | $N_{t+1} = N_t \exp(f(N_t)) + \varepsilon_t$ | $X_{t+1} = X_t + f(N_t) + \varepsilon_t$ |
| Stepwise generalized<br>logistic (SGL) | $N_{t+1} = F(1, N_t) + \varepsilon_t$ | $X_{t+1} = \ln(F(1, N_t)) + \varepsilon_t$ |

Anticipating difficulty with obtaining estimates for *θ*, we fitted the continuous-time generalized logistic (CGL) and the SGL with *θ* as a parameter but also with fixed values, ranging over the integers 1–10, 15, 20, 30, 50, 60, 100, 120, and 190.

To illustrate further the differences between the deterministic and stochastic, and between the continuous-and discrete-time, models, consider exponential growth. First note that in the absence of stochasticity,

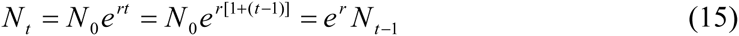

i.e., the continuous-and discrete-time models agree. Suppose there is a single stochastic perturbation to the population at time *s* < *t*. The discrete-time model incorporates this perturbation at the step following the perturbation and future projections incorporate this perturbation and are thus accurate. The naïve continuous-time model, however, continues to project future population size from the initial population size *N*_0_ and thus will differ from the actual population size at all times greater than and equal to *s*. Thus, the discrete time model will have just the one residual error when fit to actual population size while the naïve continuous-time model will have residuals for all times greater than or equal to *s*. Hence, the naïve continuous-time model can only be expected to provide a good fit to data in the absence of stochasticity or perhaps when the stochasticity in the data is small and tends to cancel out over time. In effect the naïve continuous-time model does not see stochasticity but the discrete-time model does even if *N*_*s*_ = *e*^*r*^ *N*_*s*-1_ + *e* (*e* > 0, say) but *N*_*s*+1_ = *e*^*r*^*N*_*s*_ – *d* (*d* > 0).

A more realistic approach to continuous-time dynamics with process noise was initiated by Levins (1969), who wrote *N*′ / *N = r*(*t*) with solution

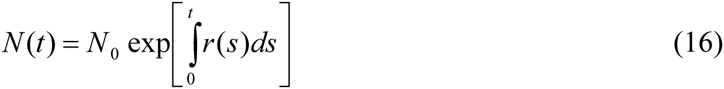

and applied the central limit theorem to the integral to deduce that it approaches a normal variate, whence *N*(*t*) is log-normally distributed. More formally, one replaces a deterministic ODE by a stochastic differential equation (SDE), in which specific parameters (e.g., *r* and/or *K*) are treated as stochastic variables (e.g., Tuckwell 1974, but there is an extensive literature on continuous-time stochastic processes and SDEs). For exponential growth, the ODE *N*′ / *N = r*, *r* constant, becomes *N*’ / *N = r*(*t*) = *r + ε* (*t*), which is formalized by the SDE

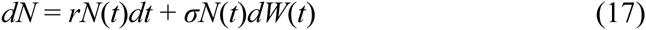

in which *ε*(*t*)*dt* has been replaced by *σdW*(*t*), with *dW*(*t*) representing the ‘differential’ of the Wiener process, representing Gaussian ‘white’ noise. For a likelihood model of the stochastic process described by an SDE, one requires the probability density *p*(*N*_*t*_,*t*,|*N*_*t-*1_) of observing abundance *N*_*t*_ at time *t* given that the abundance at time *t*-1 was *N*_*t*-1_. This probability density is obtained as the solution to the Fokker-Planck equation. When solvable, the solutions are typically analytically complicated. Various stochastic versions of the continuous-time logistic appear in the literature. Tuckwell (1974) derived *p*(*N*_*t*_,*t*,|*N*_*t-*1_) for the logistic with stochastic *r* but constant *K* but had to use Taylor series expansions to work with it. More often, for the logistic with stochastic *r* or *K*, only the steady state probability density, describing the distribution of equilibrium states, is obtained (early literature includes Levins 1969, Goel and Richter-Dyn 1974, May 1974, Karlin and Taylor 1981). The solution of the SDE (15) is known as geometric Brownian motion.

Given our data, the discrete-time models should suffice to detect density dependence if present in our data. As the SKKR population did not manifest fluctuations, more complex models of stochastic dynamics appear unnecessary for our purpose. We do note, however, that for continuous-time, stochastic exponential growth, the resulting *p*(*N*_*t*_,*t*,|*N*_*t-1*_) is log-normal and the expected abundance obeys deterministic exponential growth but with a coefficient of variation that increases exponentially as *t* → ∞ (e.g., Tuckwell 1974). The discrete-time exponential model with multiplicative error is

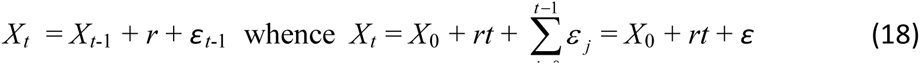

where *ε* is a sum of iid N(0,σ^2^) variates and thus N(0,σ^2^*t*). Hence, *X*_*t*_ is N(*X*_0_ + *rt*, *σ*^2^*t*), which includes the description of a single time step as ‘*X*_*t*_ is N(*X*_*t*-1_ + *r*, *σ*^2^)’. Thus the discrete-time exponential model generates the same likelihood for the data *X*_*t*_ as the model of Dennis *et al*. (1991; except that they use *μ* where we use *r* and their *r* differs from ours). Dennis *et al*. observed that these distributional assumptions for *X*_*t*_ are equivalent to *p*(*N*_*t*_,*t*,|*N*_*t-*1_) being log-normal. Hence, for a time series of abundances, the likelihood models obtained from the discrete-time exponential model with multiplicative error, from Dennis *et al*. (1991), and from the continuous-time stochastic exponential growth model are all identical, i.e., such a time series cannot distinguish these models.

Since the additive error and multiplicative error structures employ different response variables *N*_*t*_ and *X*_*t*_, respectively, distinct AIC analyses are required for these two groups of models as one cannot compare models with different response variables via AIC (Burnham and Anderson 2002:81). Since Dennis *et al*.’s model employs *N*_*t*_, rather than *X*_*t*_, as the response variable, however, its likelihood model of the time-series data can be compared via AIC_c_ to the models of Table 1 with additive error, thereby providing a common reference between the comparisons of the additive error models plus Dennis *et al*.’s model and the comparisons of the multiplicative error models since the discrete-time exponential model has the same likelihood as the Dennis *et al*. model when the latter is expressed in terms of *X*_*t*_. Thus, we consider the models listed in Table 2, together with the model of Dennis *et al*. (1991) with *N*_*t*_ as response and thus with error additive on abundance, as adequate for checking for density dependence and preferred error structure. We repeated all analyses using the time series of annual December censuses only to assess the influence of time step.

Finally, we note that the statistical assumptions regarding residuals in nonlinear regression, see below, entail that the residuals as process noise are typically interpreted as representing environmental stochasticity, which is not say that demographic stochasticity is absent from the data. Rather, more than fitting the data to such models is required to determine the nature of process noise if present in the data. We address this issue in sections 2 and 4 below

### Including additions and removals in population modelling

All introduction events were sufficiently discrete to have occurred between two consecutive semi-annual censuses. In our case there was just the one removal event and no additions occurred during the semi-annual period of Dec-06 to Jun-06 when the removals were conducted. For discrete-time models, if *n* is the net number of individuals added (with negative values of *n* accounting for a net number of removals) between *t* and *t*+1, then the model projection from *N*_*t*_ should be compared to *M*_*t*+1_ := *N*_*t*+1_ – *n* rather than *N*_*t*+1_ itself. Thus, for discrete time models, the modified census figures *M*_*t*_ were used as the response variable in the nonlinear regressions with additive error and *Y*_*t*_ := ln(*M*_*t*_) was used for the response variable for models with multiplicative error. Thus, if the deterministic model is written as *N*_*t*+1_ = *G*(*N*_*t*_), then to accommodate additions and removals we use instead *M*_*t*+1_ = *G*(*N*_*t*_).

For the continuous-time models in Table 2, projections of future abundance are made via the solution of the ODE from an initial population size. In our case, there were 5 distinct addition events, including the immigration of the one female from the other half of GFRR into SKKR, after the founding introduction in Jun-86 and one removal event. The entire period of study can be partitioned into 7 disjoint subintervals of time [0, *t*_1_], [*t*_1_, *t*_2_],…,[*t*_6_, *t*_7_] such that each subinterval consists of several consecutive between-census periods and such that each distinct addition or removal event occurred in the final between-census period of one of the first six of these subintervals. Thus, [0,*t*_1_] covers the time period from time zero to the census immediately following the first addition; [*t*_1_, *t*_2_] covers the period from *t*_1_ to the census immediately following the next distinct addition event; and so on, except that [*t*_6_, *t*_7_] covers the period from the census immediately following the final addition/removal (the removal in our case) to the final census. If *F* denotes the solution of the deterministic continuous-time model as in (3), then for the additive error models one uses: *F*(*t*, *N*_0_) to project abundance over [0, *t*_1_]; *F*(*t* – *t*_1_, *N*_*t*_1__) to project abundance over [*t*_1_, *t*_2_]; and so on, finally using *F*(*t* – *t*_6_, *N*_*t*_6__) to project abundance over [*t*_6_, *t*_7_]. These projections are compared to the modified census figures *M*_*t*_ for each census time *t*. For the models with multiplicative error, one projects abundances as just described, takes the logarithms of the projections, and compares to the *Y*_*t*_.

We also needed to modify the likelihood formulae of Dennis *et al*. (1991) to accommodate additions and removals (as they apparently did in their example of the Puerto Rican parrot, see their p. 135). Dennis *et al*. formulated a stochastic model for (st)age structured exponentially growing populations with process noise that can be fitted to time series *N*_0_, *N*_1_,…*N*_*q*_ of abundances (censuses, not estimates), with time step *τ*_*i*_ from the (*i*-1)’th observation to the *i*’th observation. The likelihood is built from the probability *p*(*N*_*i*_, *τ*_*i*_|*N*_*i*-1_) of observing *N*_*i*_ at the *i*’th observation given that the abundance was *N*_*i*-1_ at the (*i*-1)’th observation. Let *M*_*i*_ := *N*_*i*_ + removals-additions (removals and additions during the time step from the (*i*-1)’th observation to the *i*‘th observation) denote the modified count (as above). Then, in place of Dennis *et al*.’s *p*(*N*_*i*_, *τ*_*i*_|*N*_*i*-1_) we have *p*(*M*_*i*_, *τ*_*i*_|*N*_*i*-1_). In our case, the time steps are all equal. For the semi-annual censuses, *τ*_*i*_ = ½, for a time unit of one year, and *q* = 45, i.e., 45 observations (*N*_0_,*M*_1_),…,(*N*_44_, *M*_45_), each Jun and Dec from Dec-86 through Dec-08 and 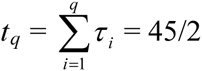. For annual censuses, *τ*_*i*_ = 1, *q* = 22, with observations (*N*_0_,*M*1),…,(*N*_21_,*M*_22_), from Dec-87 through Dec-08, and 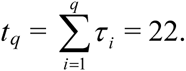.

The probability *p*(*N*_*i*_, *τ*_*i*_|*N*_*i*-1_) was derived by Dennis *et al*. from a log-normal distribution with parameters *μ* and *σ*^2^ (see their equation (8)) and the likelihood of the data (their (22)) by multiplying together such probabilities, one for each observation. Thus, to accommodate additions and removals, one literally replaces the *N*_*i*_ (but not *N*_*i*-1_) in their formula by *M*_*i*_. This substitution modifies the maximum likelihood estimates of *μ* and *σ*^2^ obtained by Dennis *et al*. as follows. In place of their equations (24) and (25), one readily obtains

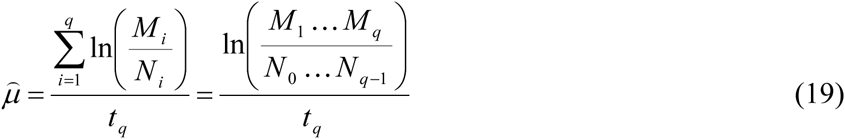

and

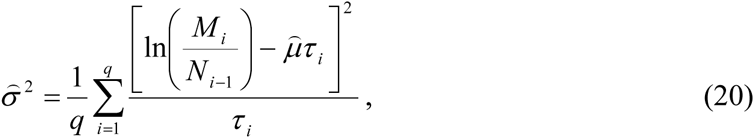

respectively, with the latter simplifying in our case to

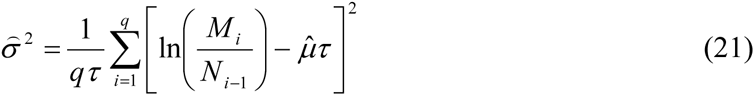

because of equal time steps. But nothing else changes for the analysis of Dennis *et al*. (1991) (though in the linear regression model one has *W*_*i*_ = ln(*M*_*i*_/*N*_*i*-1_) of course; ln(*M*_*i*_) is the modified value of ln(*N*_*i*_) and thus *W*_*i*_ still represents the increments of the Wiener process, whence the statistical properties that Dennis *et al*. appeal to remain valid and the rest of their results apply). The maximized log-likelihood is (for equal time steps)

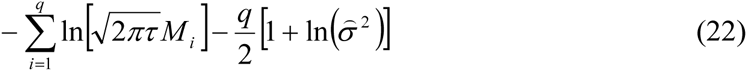

Note that, with *Y*_*t*_ = ln(*M*_*t*_), *p*(*Y*_*t*_,1|*X*_*t*_-1) is N(*X*_*t*-1_ + *r*, *σ*^2^), just as in the discrete-time exponential model with multiplicative error, so the equivalence of the likelihood descriptions of a time series of abundances is maintained when introductions and removals are accounted for.

### Nonlinear regression and AIC_c_

For a model of the form

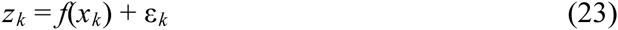

with ε_k_ iid N(0,σ^2^) and *n* observations, the log-likelihood is

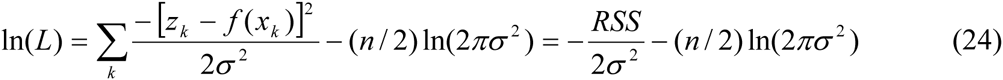

where *RSS* is the residual sum of squares (Burnham and Anderson 2002:12, 108–109; Bates and Watts 1988:4). Maximum likelihood (ML) estimation of the structural parameters in *f* is equivalent to (non-linear) least-squares estimation. ML estimation of σ^2^ is found easily by calculus to be

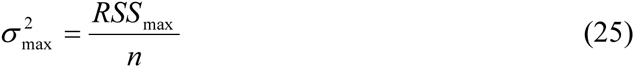

where *RSS*_max_ denotes the residual sum of squares evaluated for the ML estimates of the structural parameters of *f* (i.e., what is usually meant by the residual sum of squares). The deviance (i.e., -2 times the maximized log-likelihood) is then found to be

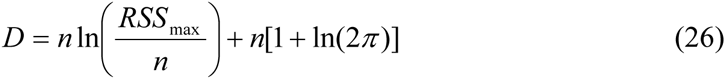

Nonlinear regression was performed in the nonlinear estimation module of Statistica 8 (Statsoft), which provided the ML estimates of the structural parameters of the model and the residuals. From the residuals we computed *σ*^2^_max_ and the deviance *D*. AIC_c_ was then computed as

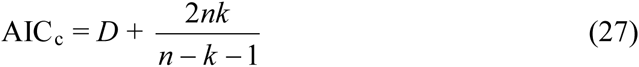

(Burnham and Anderson 2002), where *k* is the number of estimated parameters, here the number of structural parameters in the model plus one (for *σ*^2^), and *n* is the number of data points used in the likelihood. Note that a time series of length one does not permit estimation for any of the models in Table 1, i.e., the initial population census does not count as a datum point in the likelihood. As noted above in the discussion of Dennis *et al*.’s model, the 23 years of Jun and Dec censuses yield 45 semi-annual census data of the form (*M*_*t*+1_, *N*_*t*_) and 22 annual census data of a similar form. Thus, for all models, *n* = 45 for the semi-annual census data and 22 for the annual census data.

For AIC_c_ calculations, the second term in (26) is common to all nonlinear regression models so cancels out in computations of ΔAIC_c_ for such models. However, for comparisons of the models with additive error with the model of Dennis *et al*. (1991) it is essential to retain that term in the deviance and all AIC_c_ calculations.

Statistica provides an *R*^2^ value for each nonlinear regression, computed as follows. The total sum of squares (SS) is defined as usual for a response variable *z* as 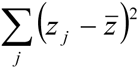 and the Error SS is defined as 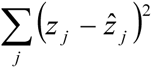, where 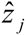 is the predicted value. Statistica then defines the regression SS as Total SS – Error SS and *R*^2^ as the ratio of regression SS to total SS, i.e., as 1 – (the ratio of Error SS to Total SS), as a measure of variation explained by the model.

Nonlinear regression requires starting values (SV) for the structural parameters to be estimated (Bates and Watts 1988). For *r* we used 0.05 as SV for both semi-annual and annual census analyses, which proved unproblematic. For *K* we began with an SV of 150. Since the dataset gave no indication that an equilibrium population size had been reached, we expect *K* to be larger than 110 (mean density 0.5 rhino/sq. km), the Dec-08 census figure. The highest local density for black rhinoceros reported by Owen-Smith (1988:224) was 1.6 rhino/sq. kms. A mean density of 1.6 should therefore provide an upper bound on *K*, yielding 352. In fitting the generalized logistic with *θ* fixed, we found that we had to lower the SV of *K* as *θ* increased in order for Statistica to fit the model (otherwise it complained that the remaining model parameters were ‘probably very redundant; estimates suspect’). Larger SVs for *K* did not help.

For models in which *θ* was to be estimated in our analyses, we tried SVs of 1 and 4.5 (the latter following Eberhardt *et al*. 2008). For the semi-annual censuses, for CGL with additive error, when SV was set to one, Statistica reported model parameters were ‘probably very redundant; estimates suspect’, but fit the model to the data with an SV of 4.5. The opposite was the case for the CGL with multiplicative error. The resulting estimates of *θ* of 11.3 and 1.3, respectively, may reflect a dependence on SVs due to the redundancy. Both estimates had high CV, 3.6 and 0.9, respectively. (In the case of multiplicative error, the estimate of *K* was essentially the same estimate as obtained for CGL with *θ* fixed at one.) For the DGL, genRicker, and SGL, estimates were obtained with both SVs for both error structures. An SV of 1 for *θ* either yielded the same result as an SV of 4.5, or an estimate for *K* greater than 1000, which is unrealistically high, and an estimate of *θ* less than 1.5. On the other hand, an SV of 4.5 for *θ*, yielded estimates of *K* less than 200 and estimates of *θ* between 5 and 10, with each estimate similar across models. But in all cases, CVs of estimates of *K* and *θ* were very high, greater than 10, making the estimates uninformative. For the annual censuses, similar dependence on SVs was observed for additive error models, but for multiplicative error both SVs of 1 and 4.5 returned similar estimates of *K* > 1000 and *θ* < 1. Again CVs were larger than 10. Given that our dataset turned out to be well modelled by exponential growth, the various versions of generalized logistic (CGL, DGL, genRicker, SGL) approximate the exponential with either large values of *K* or large values of *θ*; which results in considerable redundancy between *K* and *θ* for such data. Statistica, as noted, complained about such redundancy. When estimates were obtained, their CVs indicated these estimates were of no value. Thus, our general conclusion about this data set (exponential growth with no information on *K* or *θ*) does not depend on SVs.

For large herbivores, one expects *θ* > 1 (Owen-Smith 2010). This expectation appears to be challenged by Sibly *et al*. (2005), who fitted numerous time series of abundances from the Global Population Dynamics Database (GPDD; http://www3.imperial.ac.uk/cpb/databases/gpdd) to the generalized Ricker model with multiplicative error and obtained more often than not values of *θ* < 1 and even negative. For critical assessments of these analyses and Sibly *et al*.’s response, see *Science* **311** (2006), p.1100d, and further see Doncaster (2008), Eberhardt *et al*. (2008), Polansky *et al*. (2009), and Clark *et al*. (2010). The values of *θ* obtained by Sibly *et al*. are appended to the corresponding time series in the GPDD. We inspected all time series for Rhinocerotidae, Elephantidae, Giraffidae, Hippopotamidae, Bovidae, and Cervidae and found little evidence to contradict the expectation of *θ* > 1 for large herbivores. There were actually few such time series for which *θ* was estimated and for those time series from robust studies *θ* was estimated to be greater than one, except in the one case of Owen-Smith’s (1990) study. But Owen-Smith found that fluctuations in abundance in that study were significantly influenced by exogenous factors in addition to density, which may have complicated estimates of *θ*.

### Results of scalar population model comparisons: semi-annual time step

**Table 3.** Results of nonlinear regression of scalar models in Table 2 for the semi-annual census data. Model name abbreviations as in table 2; *R*^2^ is 1 – (ratio of error sum of squares to total sum of squares) for the regression fit; Dev is the deviance, -2 times the maximized log-likelihood; *k* is the number of estimable model parameters (including the variance of the residuals); *r* ± SE is the estimated value of the parameter *r* common to all the models as an *annual* rate ± its SE; ΔAIC_c_ is the model’s AIC_c_ value minus that of the model with the smallest AIC_c_ value, which was the discrete-time exponential model for both error structures.

| Model | Additive error |  |  |  |  | Multiplicative error |  |  |  |  |
| --- | --- | --- | --- | --- | --- | --- | --- | --- | --- | --- |
| | $R^2$ | Dev | $k$ | $r \pm \text{SE}$ | $\Delta\text{AIC}_c$ | $R^2$ | Dev | $k$ | $r \pm \text{SE}$ | $\Delta\text{AIC}_c$ |
| Cexp | 0.996 | 194.9 | 2 | 0.0912<br>$\pm 0.0023$ | 25.6 | 0.995 | -99.4 | 2 | 0.1016<br>$\pm 0.0043$ | 30.4 |
| CGL | 0.996 | 194.1 | 4 | 0.0921<br>$\pm 0.0027$ | 29.5 | 0.996 | -106.3 | 4 | 0.115<br>$\pm 0.011$ | 28.3 |
| Dexp | 0.998 | 169.3 | 2 | 0.1017<br>$\pm 0.0092$ | 0 | 0.997 | -129.9 | 2 | 0.102<br>$\pm 0.017$ | 0 |
| DGL | 0.998 | 169.3 | 4 | 0.106<br>$\pm 0.021$ | 4.7 | 0.997 | -129.9 | 4 | 0.105<br>$\pm 0.021$ | 4.7 |
| genRicker | 0.998 | 169.3 | 4 | 0.103<br>$\pm 0.020$ | 4.7 | 0.997 | -129.9 | 4 | 0.102<br>$\pm 0.020$ | 4.7 |
| SGL | 0.998 | 169.3 | 4 | 0.103<br>$\pm 0.020$ | 4.7 | 0.998 | -130.7 | 4 | 0.102<br>$\pm 0.020$ | 3.9 |

Comparisons of the models in Table 2 for the semi-annual censuses are presented in Table 3. The discrete-time exponential was unambiguously the best model of the data for both error structures. While estimates of *r* were relatively consistent across models and precise, estimates of *K* and *θ* were not. For the CGL, *K* was estimated as 127 (additive error) and 196 (multiplicative error) each with CV = 0.8, while DGL, genRicker, and SGL gave estimates near 190 (additive error) and near 169 (multiplicative error) but CVs exceeded 10. For the CGL *θ* was estimated as 11.3 (additive error) with CV = 3.6 and 1.3 (multiplicative error) with CV = 0.9, and more consistently across DGL, genRicker, and SDL at about 5.5 (additive error) and 7.5 (multiplicative error), but with CVs exceeding 10.

For additive error, the deviance of the CGL models with fixed *θ* increased slightly from 191.9 for *θ* = 1 to 194.4 for *θ* = 4, then decreased to 194.1 for *θ* = 10, and then increased monotonically to 194.9 for *θ* = 190 while for multiplicative error the deviance increased monotonically from a low of -106.0 for *θ* = 1 to -99.6 for *θ* = 190. Estimates of *K* for CGL with fixed *θ* and additive error decreased from 545 with CV = 0.5 (*θ* = 1) to 110 with CV = 0.03 (*θ* =190) with a similar pattern for multiplicative error beginning with an estimate of 259 with CV = 0.3 for *θ* = 1 and a CV of 0.08 for *θ* = 190. For the SGL with fixed *θ*, for both error types, deviance did not vary with *θ*, but while estimates of *K* roughly decreased to near 110 with increasing *θ*, their CVs did not and were consistently much greater than one.

Differences in ΔAIC_c_-values amongst the discrete-time models in Table 2 were due almost entirely to the number of model parameters. The CGL models with fixed *θ* had ΔAIC_c_ similar to the other continuous-time models and so were not at all competitive. The SGL models with fixed *θ*, having only slightly larger deviance than the discrete-time exponential and only one more parameter were competitive with ΔAIC_c_ values of about 2.3 (additive error) and 1.5 (multiplicative error) but did not, as noted above, yield informative estimates of *K*.

### Results of scalar model comparisons: annual (December) Censuses

As for the semi-annual censuses, for the annual census data the discrete-time models exhibited the same deviances so that their ΔAIC_c_ values differed according to their number (*k*) of model parameters. Unlike the semi-annual censuses, however, the (naïve) continuous-time models exhibited lower deviances and lower AIC_c_ values than the discrete-time models. For both error types, the continuous-time exponential model had lowest AIC_c_ amongst the models in Table 2, though the CGL had lower deviance and in the case of multiplicative error the difference in AIC values between the two continuous-time models was marginal.

For CGL, the estimate of *K* was 117 with CV = 0.4 (additive error) and 382 with CV = 2.1 (multiplicative error), while for DGL, genRicker, and SGL about 150 with CV about 6 (additive) and over 1000 with CV about 35 (multiplicative). For CGL, the estimate of *θ* was about 16 with CV about 4 (additive error) and 0.7 with CV = 1.6 (multiplicative error), while for DGL, genRicker and SGL about 7.5 with CV about 15.5 (additive error) and 0.8 with CV about 14 (multiplicative error).

**Table 4.** Results for the comparison of scalar models of Table 2 for the December census data only. Model name abbreviations as in Table 2; *R*^2^ is 1 – (ratio of error sum of squares to total sum of squares) for the regression fit; Dev is the deviance, -2 times the maximized log-likelihood; *k* is the number of estimable model parameters (including the variance of the residuals); *r* ± SE is the estimated value of the parameter *r* common to all the models as an *annual* rate ± SE; ΔAIC_c_ is the model’s AIC_c_ value minus that of the model with the smallest AIC_c_ value, which was the continuous-time exponential (Cexp) for both analyses.

| Model | Additive error |  |  |  |  | Multiplicative error |  |  |  |  |
| --- | --- | --- | --- | --- | --- | --- | --- | --- | --- | --- |
| | $R^2$ | Dev | $k$ | $r \pm \text{SE}$ | $\Delta\text{AIC}_c$ | $R^2$ | Dev | $k$ | $r \pm \text{SE}$ | $\Delta\text{AIC}_c$ |
| Cexp | 0.995 | 100.0 | 2 | 0.0928 | 0 | 0.996 | -57.8 | 2 | 0.1005 | 0 |
| | | | | $\pm 0.0033$ | | | | | $\pm 0.0049$ | |
| CGL | 0.995 | 98.8 | 4 | 0.0940 | 4.5 | 0.997 | -63.1 | 4 | 0.123 | 0.5 |
| | | | | $\pm 0.0037$ | | | | | $\pm 0.029$ | |
| Dexp | 0.994 | 105.8 | 2 | 0.101 | 5.8 | 0.995 | -52.2 | 2 | 0.104 | 5.6 |
| | | | | $\pm 0.011$ | | | | | $\pm 0.016$ | |
| DGL | 0.994 | 105.8 | 4 | 0.109 | 11.5 | 0.995 | -52.3 | 4 | 0.117 | 11.3 |
| | | | | $\pm 0.024$ | | | | | $\pm 0.079$ | |
| genRicker | 0.994 | 105.8 | 4 | 0.103 | 11.5 | 0.995 | -52.3 | 4 | 0.110 | 11.3 |
| | | | | $\pm 0.021$ | | | | | $\pm 0.070$ | |
| SGL | 0.994 | 105.8 | 4 | 0.103 | 11.5 | 0.995 | -52.5 | 4 | 0.110 | 11.3 |
| | | | | $\pm 0.021$ | | | | | $\pm 0.068$ | |

For additive error, the deviance of the CGL models with fixed *θ* decreased (monotonically with increasing *θ* from *θ* = 1 to *θ* =190) slightly from 99.8 to 99.2 so ΔAIC_c_ decreased from 2.5 to 1.9. The estimates of *K* decreased monotonically from 1380 with a CV of 2.2 (*θ* = 1) to 110 with CV = 0.2 (*θ* = 190). For multiplicative error, deviance *increased* from -62.9 (*θ* = 1) to -58.1 (*θ* = 190) so that ΔAIC_c_ (relative to the continuous-time exponential) increased from -2.4 to 4.8. Estimates of *K* followed a similar pattern as for additive error. Thus, the (naïve) continuous-time logistic (i.e., *θ* = 1 in the CGL) actually gave the best fit to the data for multiplicative error (Δ AIC_c_ = -2.4 relative to the continuous-time exponential). The estimate of *K* was 270±100, i.e., CV = 0.37. While this range is biologically plausible, the behaviour of the estimates of *K* as *θ* increased indicated that the nonlinear regression was estimating *K* so that the inflexion point (5) of the solution lay beyond the observed data. For additive error, increasing *θ* slightly improved the fit, i.e., these models became more competitive as they better approximated threshold-like models with exponential growth for the actual data, but the opposite was true for multiplicative error, unlike for the semi-annual census data. The fact that the naïve continuous-time models were favoured in the analysis of the annual census data suggests that the stochasticity in the semi-annual census data tended to average out over the annual time step. For the SGL models with fixed *θ*, for both error types, deviances and AIC_c_ values did not vary to any important degree and were not competitive (ΔAIC_c_ greater than eight for both error types). Estimates of *K* started unrealistically high with large CVs for *θ* = 1, decreased monotonically over the rang *θ* = 1 to 10 to between 150 and 110 with CVs of about 0.7 but for higher values of *θ* the estimates of *K* fluctuated outside that range and had extremely large CVs.

Figure 2 shows a plot of the pairs (*X*_*t*-1_, *Y*_*t*_) for the semi-annual census data, i.e., of the log-transformed census data, modified to account for additions/removals, as the second coordinate (*Y*_*t*_ = ln(ModCount)) versus the log-transformed actual census data at the previous time as the first coordinate (*X*_*t*-1_ = ln(PrevCount)), together with the line *y* = x + *c*, where *c* is the estimate of (the semi-annual rate) *r* from the discrete-time exponential model with multiplicative error, i.e., the overall best fit model. Thus, the line represents this model, viz., *Y*_*t*_ = *X*_*t*-1_ + *r* + *ε*_*t*_. Figure 3 shows the same plot for the annual census data with *c* the estimate of (the annual rate) *r* from the discrete-time exponential model with multiplicative error fit to that data. Note that we have not used the jitter option to separate data that coincide, as that would defeat comparison of the fit to the line. In Figure 2, for example, for the first four data (for Dec-86, Jun-87, Dec-87, Jun-88, the *Y*_*t*_ and *X*_*t*-1_ values are equal and so plot as the same point, below the line (at *X*_*t*-1_ = 1.1). Though visual inspection does not quantify the model fit as well as the deviances of the models, note that beyond 3.5 on the horizontal scale in Figure 3, there appears to be a tendency for the data to fall just below the line. It is not just that the June census data has been removed from the plot in Figure 2, but the remaining (December) data is now fit to the exponential model by projecting that data over the larger time step, i.e., whereas *Y*_*t*_ is projected from *X*_*t*-1_ in Figure S2, it is projected from *X*_*t*-2_ (using the same parametrization of censuses as for the semi-annual data) in Figure 3, resulting in the different estimate of *r* (as an annual rate).

**Figure 2.**
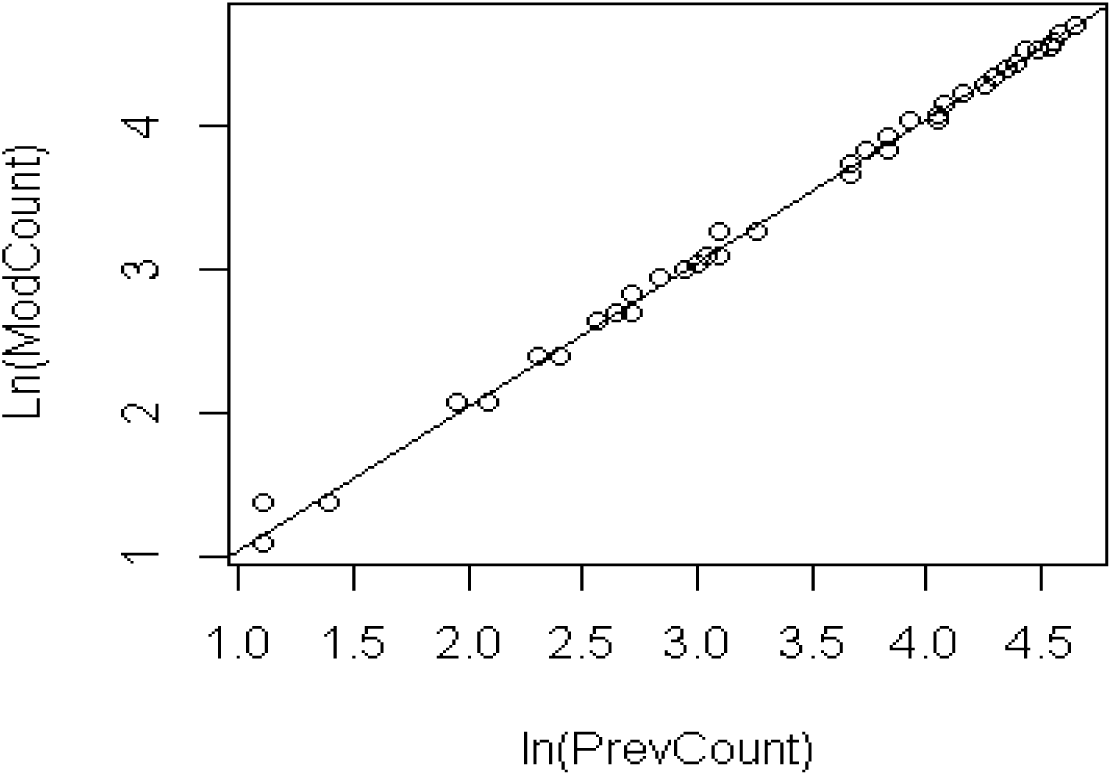
The semi-annual census data plotted together with the line *y* = *x* + *c*, *c* = the estimate of *r* from the discrete-time exponential model with multiplicative error fit to that data.

**Figure 3.**
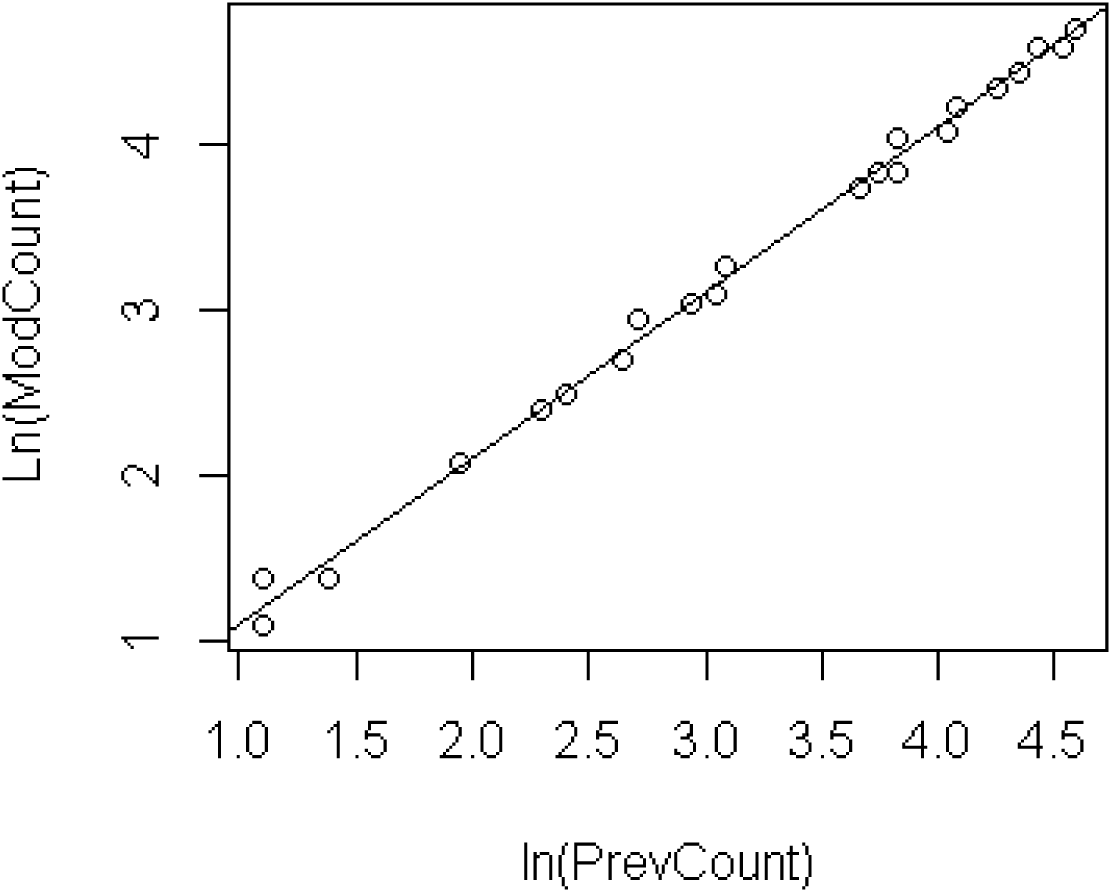
The annual census data plotted together with the line *y* = *x* + *c*, *c* = the estimate of *r* from the discrete-time exponential model with multiplicative error fit to that data.

Similarly, the differences between the model fits of the continuous-time exponential and logistic (i.e., *θ* = 1 in the CGL) and the discrete-time exponential for annual census data and multiplicative error structure appears fairly subtle in a graphical display (Figure 4). The sum of squared residuals for the continuous-time logistic was 0.0733 versus 0.0931 for the continuous-time exponential, the squared residuals of the latter consistently slightly larger than those of the former for years 7 – 17, summing to 137% of the difference in the sum of squared residuals for these two models. Thus, the lower deviance for the continuous-time logistic, which yields a lower AIC_c_ by 2.4 units, despite the extra model parameter, reflects better model fit for years 7 – 17 rather than indicating a slowing of population growth rate towards the end of the study. The sum of squared residuals of the discrete-time exponential was 0.1200, about 1.6 times larger than that for the continuous-time logistic, which translated into some 8 AIC_c_ units difference. Thus, there is no evidence that the model fits to the longer time-step are more informative than the fits to the semi-annual census data. That smaller residuals occurred for the fit of a continuous-time model than for a discrete-time model indicated that the annual census data was more easily fit with a projection from an initial value rather than an adjustment each time step, i.e., that the annual census data smoothed out the irregularities in the semi-annual census data.

### Dennis *et al*. (1991) model

The computation of Dennis *et al*.’s estimate of their *μ* from (19) agreed to nine decimal places with the Statistica estimates of *r* for the discrete-time exponential model using either semi-annual or annual census data. Agreement of estimates of SEs was less close, to eight decimal places using the annual census data and seven using the semi-annual census data. Agreement for the estimates of *σ*^2^ was to more than 10 decimal places using either set of data. For both semi-annual and annual census data, the Dennis *et al*. model had a lower deviance than any model with additive error. For semi-annual census data it was almost 10 AIC_c_ units below the discrete-time exponential model and thus was unambiguously the best model amongst those models. For the annual census dataset, the Dennis *et al*. model was about 7.7 AIC_c_ units below the (naïve) continuous-time exponential model and again unambiguously the best model amongst those models. Identifying the Dennis *et al*. model with the discrete-time exponential model with multiplicative error structure, then for semi-annual census data, one concludes that model is overall the best fit to the data, even across error structures. This conclusion fails for the annual census data, as (naïve) continuous time exponential and logistic models with multiplicative error outcompeted the discrete-time exponential model with multiplicative error structure.

**Figure 4.**
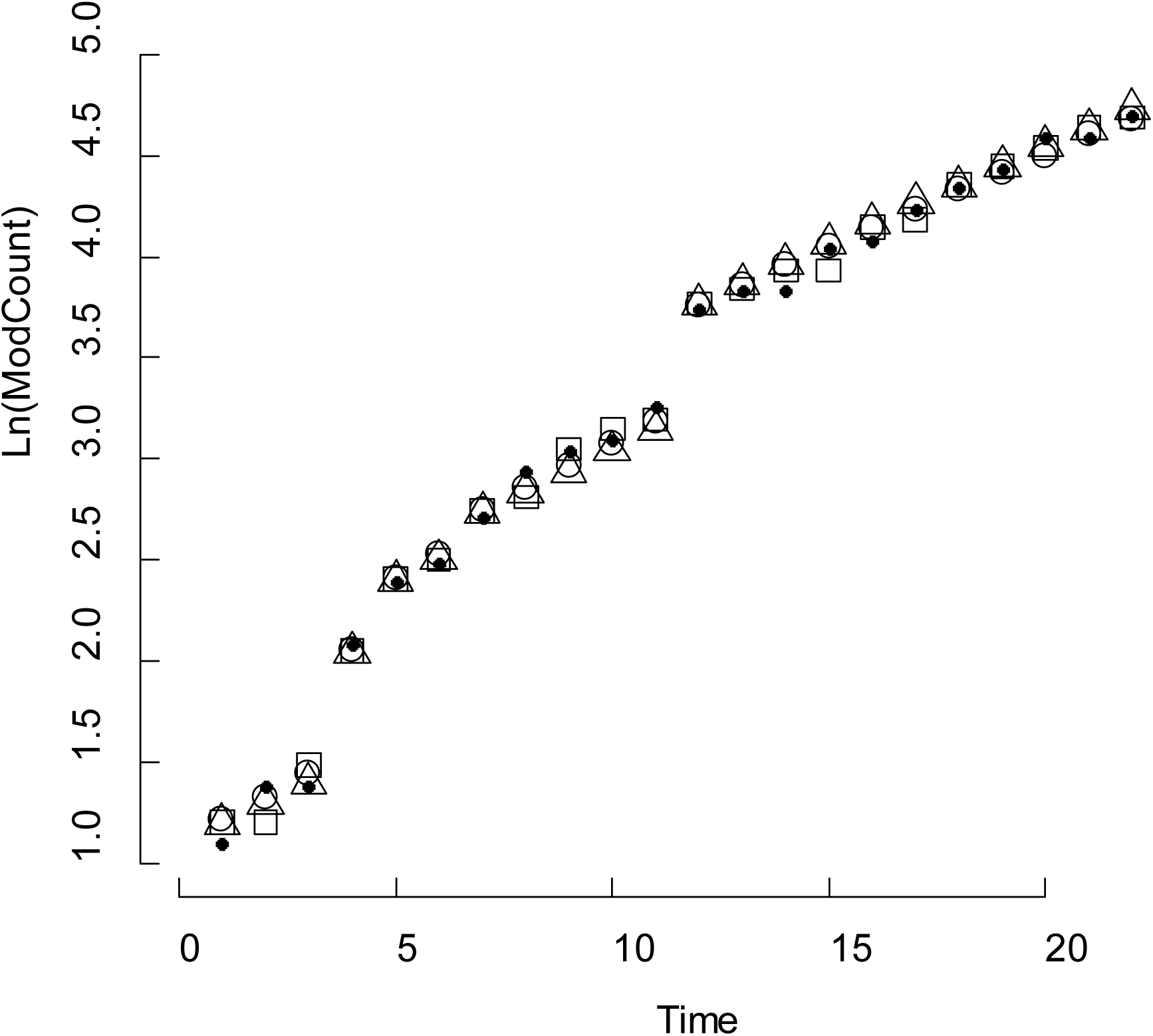
Annual census data, multiplicative error structure. Response variables plotted on the log scale: observed (modified) count (∂) predicted responses for the continuous-time exponential (Δ), continuous-time logistic (○), and discrete-time exponential (□).

## 2. Estimating Demographic and Environmental Stochasticity from Scalar Models

Process noise, i.e., departure from the deterministic model, is interpreted as arising from demographic and environmental sources of stochasticity. For a population of statistically identical individuals, Engen *et al*. (1998) provided a decomposition of the variance in the change in population size from a given population size, which provided definitions of demographic and environmental stochasticity and demographic covariance. Sæther *et al*. (1998a) showed how to apply this formalism to demographic data and a time series of abundances (outlined in Lande *et al*. 2003; see also Morris and Doak 2002:127–133).

Environmental stochasticity is typically construed as variations from year to year rather than at finer scales, so we conducted this analysis on an annual basis. For each end-of-year (i.e., our December censuses), 1986 through 2007, we tabulated the contribution each individual alive at that time made to the following end-of-year census: counting one for survival and one for producing an offspring during that year that survived to the end of the year. The usual estimate of variance applied to this annual data defines a quantity *V*_*d*_(*N*) for each such end-of-year, which is parametrized by the population size rather than time. As noted above, the formalism assumes that individuals are statistically identical; in particular, for any given year, the individuals alive at that time are assumed to have the same expected contribution to the population the following year. This assumption thus ignores, for example, stage differences such as the difference between immature individuals that contribute only by survival and mature individuals that can also contribute by reproduction. Typically, the formalism is applied to subunits of a population (e.g., females) that can plausibly be treated as homogeneous and to populations in which individuals mature over one time step. Since we are attempting to interpret the process noise of a scalar population model, which also neglects differences between individuals, we proceed as if the formalism is applicable to our data noting that our estimate of *V*_*d*_(*N*) conflates strict demographic stochasticity and fixed demographic differences between individuals (such as stage differences), but is still demographic in nature. This conflation may lead to biases in estimates of probability of extinction and time to extinction (Fox and Kendall 2002, Kendall and Fox 2002, Morris and Doak 2002:132–133, Melbourne and Hastings 2008) but our concern is only to estimate the relative demographic and environmental contributions to process noise.

As the best description of the semi-annual data was the discrete-time exponential model, we took that model as the best deterministic model for the SKKR population dynamics and applied it to the December annual census data, i.e., we converted the value of *r* obtained from the discrete-time exponential model with multiplicative error for the semi-annual census data, to an annual time step. The prescription in Sæther *et al*. (1998a) (also Lande *et al*. 2003, equation (1.11); Morris and Doak 2002, equation (4.14)) amounts to putting

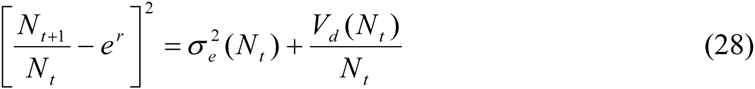

where *N*_*t*_ is the observed abundance at time *t* (the December for which one computes the contributions of living individuals to time *t*+1) and *N*_*t*+1_ is the abundance at time *t*+1. For our data, we must replace *N*_*t*+1_ by *M*_*t*+1_, the modified count at *t*+1 to account for any introductions or removals during the time step (see **Including additions and removals in population modelling** above). Using the estimate of *V*_*d*_(*N*_*t*_) obtained as described in the previous paragraph, one obtains from (28) an estimate *σ*_*e*_^2^(*N*_*t*_) of the environmental contribution to process noise.

The advice (Sæther *et al*. 1998a, Morris and Doak 2002, Sæther and Engen 2002, Lande *et al*. 2003) is then to regress each of *V*_*d*_ (*N*_*t*_) and *σ*_*e*_^2^(*N*_*t*_) on *N*_*t*_ to check for density dependence. For *V*_*d*_ (*N*_*t*_), regression yielded a slope of -0.00040 (*p =* 0.55) and with a positive intercept (*p* = 0.00016), while for *σ*_*e*_^2^(*N*_*t*_), regression yielded *p* > 0.2 for both slope and intercept. The *t*-test for each set of *V*_*d*_ (*N*_*t*_) and *σ*_*e*_^2^(*N*_*t*_) values with a null hypothesis of zero mean returned *p* = 0.000001 for the former (mean = 0.137 ± 0.091) and *p* = 0.44 for the latter. Thus, density dependence was detected for neither *V*_*d*_ (*N*_*t*_) nor *σ*_*e*_^2^(*N*_*t*_) but the mean of *V*_*d*_ (*N*_*t*_) is judged to be nonzero. Overall estimates of the demographic and environmental contributions to process noise are obtained as weighted means of *V*_*d*_ (*N*_*t*_) and *σ*_*e*_^2^(*N*_*t*_) (Sæther and Engen 2002:194, 197), for which we obtained 0.127 and 0.0002, respectively. The estimate of the demographic component can be decomposed into survival and fecundity components and the covariance between these two components; for each computation of *V*_*d*_(*N*_*t*_) one separates those contributions that are due to survival from those due to reproduction. The weighted means were 0.109 for the fecundity component and 0.015 for the survival component. Since only 14 mortalities contributed to the survival component, it is not surprising that the fecundity component was the more important contribution (86%).

Our strategy of including all individuals for the computation of demographic stochasticity is conservative; it is more usual to restrict to the female segment of the population. Doing so returned an estimate of 0.178, in place of 0.127, for σ_*d*_^2^, without altering any other conclusions of the previous paragraph other than increasing the component of σ_*d*_^2^ due to fecundity to 92%.

## 3. Matrix Model

All matrix computations were performed in R 2.15.1(R Development Core Team. 2009. R: A language and environment for statistical computing. R Foundation for Statistical Computing, Vienna, Austria. ISBN 3-900051-07-0, URL http://www.R-project.org).

Since the semi-annual census data was best modelled by the discrete-time exponential, we constructed a stage-based, two-sex, birth-flow matrix model with semi-annual time step using the entire population history from Jun-86 to Dec-08. As a result estimates are not based on samples and there are no ranges or SEs for parameter estimates. Consequently, quantities computed from the matrix models, e.g., *λ*, are exact for our data, i.e., do not possess sampling distributions or SEs. We used a semi-annual time step for greater accuracy of modelling dynamics by matrix model projections and thus our time unit in all computations is half a year. We employed a stage-based rather than age-based model as life stages defined as biological states are more relevant than age (Law and Linklater 2014). For each sex, the biological states of interest are calf (C), subadult (S), and adult (A) (prefixed by F or M to specify sex, e.g., FC for female calf), as defined in Table 1.

Lacking paternity data, we employed Goodman’s (1969) two-sex model in which reproduction is attributed to females only (see also Charlesworth 1994:6–7). The projection matrix *A* for Goodman’s model takes the form

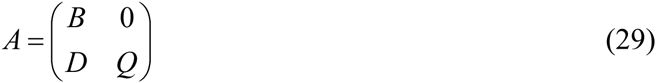

where the matrix *B* is the female-only matrix model for the population dynamics, the matrix *Q* is the analogue for males but encodes only survival as males are not modelled as contributing to reproduction, and the matrix *D* encodes the production of male offspring by females. The matrix *A* is reducible (Caswell 2001:90) but, assuming *B* and *Q* are irreducible, for a realistic two-sex model, i.e., one in which the stable-stage distribution (SSD) contains both males and females, one can show that the dominant eigenvalue *λ*_*A*_ of *A* has (right) eigenvector (*w*_*F*_,*w*_*M*_) with each of *w*_*F*_ and *w*_*M*_ nonzero and with strictly positive components. Moreover: *λ*_*A*_ is the dominant eigenvalue of *B* with (right) eigenvector *w*_*F*_; the left eigenvector of *A* (i.e., the reproductive value vector when appropriately scaled) is (*v*_*F*_,**0**) where *v*_*F*_ is the left eigenvector of *λ*_*A*_ as dominant eigenvalue of *B*; the sensitivities of *λ*_*A*_ with respect to entries of *D* and *Q* are zero and both the sensitivities and elasticities of *λ*_*A*_ with respect to entries of *B* are unambiguous as to whether one considers them as properties of the two-sex or female-only model.

The nonzero entries of matrix *B* consist of transition rates *G* between stages, survival rates *P* within stages, and fecundity rates *F* for the production of female offspring. The matrix *Q* will have an identical structure except that where the fecundity rates occur in *B* the corresponding entries in *Q* are zero. The only nonzero entries of *D* are for the fecundity rates for the production of male offspring from females. Though the stages C, S, and A are of primary biological interest, for a semi-annual time step we could build a more accurate matrix model for SKKR as regards transition rates by partitioning C and S into substages. For SKKR, calves became subadults beyond 1.5 years of age and subadults became adults beyond 2.5 years of having become a subadult. There were calves of both sexes that did become subadults before the age of two and females that became adults in their third year of being a subadult. As males were not considered adult until age eight, they became adult at least a year later than females became adults.The purpose of the substages was to exclude transitions from the first year as a calf and the first two years as a subadult. The ‘stages’ for the matrix model then were C1a (calf at most 6 months old), C1b (calf, 6 months < age ≤ 12 months), C2 (calf, age > 12 months), S1a (subadult, within 6 months of becoming subadult), S1b (subadult, time since becoming subadult greater than 6 months but less than or equal to 12 months), S2a (subadult, time since becoming subadult greater than 12 months but less than or equal to 18 months), S2b (subadult, time since becoming subadult greater than 18 months but less than or equal to 24 months), S3 (subadult, time since becoming subadult greater than24 months), A (adult), for each sex. So as to retain the term ‘stage’ for the biological states of C, S, A, we refer to these ‘stages’ as substages. The matrix *B* took the form

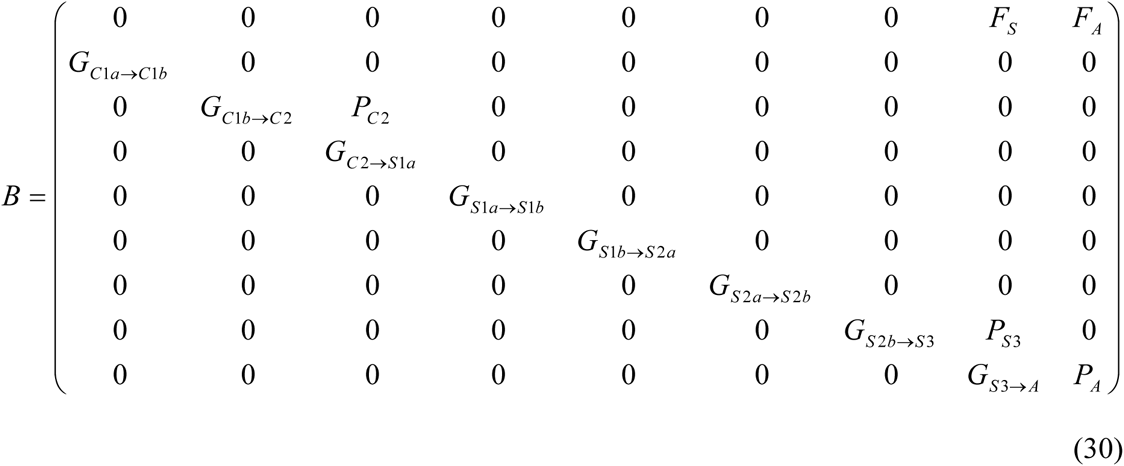

As noted already, the matrix *Q* takes the same form except that the *F*_*S*_ and *F*_*A*_ entries are zero, while the matrix *D* has entries *M*_*S*_ and *M*_*A*_ in the final two columns of its first row (for the production of male offspring) as its only nonzero entries. We parametrized these matrices, i.e., estimated their nonzero entries, in two different ways, yielding two realizations of the matrix model. All matrix-model analyses were performed with both parametrizations as a check on the robustness of results.

### MM1 matrix model parametrization

For the first parametrization, denoted MM1, *P*’s and *G*’s were modelled in terms of probability of transition between stages and survival during stages. For each sex and stage (not substage), we computed the ratio of the number of individuals of that sex that died during that stage to the total time individuals of that sex and stage were at risk (i.e., alive) as an estimate of mortality rate and subtracted this quantity from one to obtain a sex-specific, stage-based survival rate *σ* (e.g., Brault and Caswell 1993). In this parametrization we did not distinguish survival for substages of a given stage because we regard stage as the state of biological interest and substages as conveniences for model parametrization. For each of C1a → C1b, C1b → C2, S1a → S1b, S1b → S2a, S2a → S2b, S2b → S3, transition is automatic given survival over the time step. Thus, for each sex,

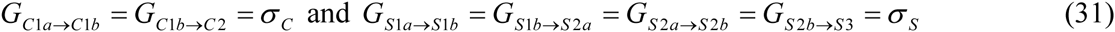

The probabilities of transitions C2 → S1a and S3 → A were estimated as follows. For FC2 → FS1a, we computed the mean duration of calfhood for those females that were born and transitioned from calf to subadult during the study, subtracted 1 year (i.e., two time units) from this mean, and took the reciprocal to define the probability *γ*_*FC2*_ →_*FS*1*a*_ (Brault and Caswell 1993, Caswell 2001, §6.4.1). For FS3 → A, we computed the mean duration of subadulthood for those females that transitioned from calf to subadult and subadulthood to adulthood during the study, subtracted 2 years (i.e., four time units) from this mean, and took the reciprocal to define the probability *γ*_*FS# → FA*_. Analogous quantities were computed for the transitions MC2 → MS1a and MS3 → MA. Now, suppose there are *n*(*t*) individuals in some specific substage (FC2, MC2, FS3, or MS3) at time *t*. In reality, transitions to the next substage (FS1a, MS1a, FA, MA, respectively) can occur at any time between *t* and *t*+1. Let *u* be an element of [0,1]. If all transitions occur at time *t*+*u*, with probability *γ* and if *ς* is the survival rate for the stage that individuals transition from and *σ* is the survival rate of the stage to which individuals transition to, then

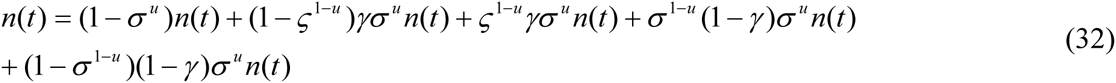

i.e., the *n*(*t*) can be written as the sum of : those that don’t survive until time *t*+*u*; those that survive until *t*+*u*, transition to the next stage but don’t survive until time *t*+1; those that survive until *t*+*u*, transition to the next stage and survive until time *t*+1; those that survive until time *t*+*u*, don’t transition to the next stage and survive until time *t*+1; and those that survive until time *t*+*u*, don’t transition to the next stage and don’t survive until time *t*+1.

Hence, the transition rate from the one stage to the next and the persistence rate within the initial stage are, respectively

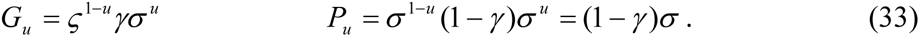

Taking the mean over *u* in [0, 1] yields

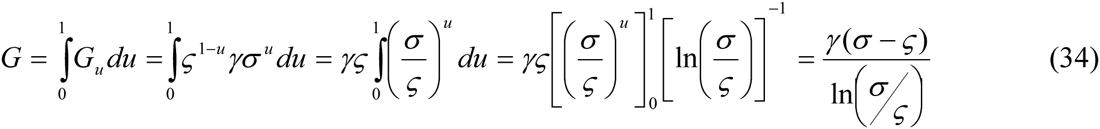

and

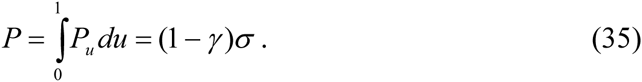

We used formula (34) for *G*_*C*2→*S* 3_ and *G*_*S* 3→ *A*_ and (35) for *P*_*C* 2→*C* 2_, *P*_*S* 3→*S* 3_, *P*_*A*→ *A*_, for each sex.

Reproduction takes place either by existing female adults or by female subadults that transition to adulthood by virtue of giving birth. For fecundity *F* we first computed the fertility (i.e., birth rate) *m* of adult females as the ratio of the number of births (of a specific sex), excluding births that initiated the transition of the mother from subadulthood to adulthood, to the total number of female-adult time units during the study. We computed the probability *α* that a female transitioned from subadulthood to adulthood by giving birth (as opposed to reaching the age of seven years without having given birth, see Table 1) as the ratio of number of females that did so transition to the total number of females that transitioned from subadulthood to adulthood during the study. Let *φ* be the birth sex ratio for females, i.e., F/(M+F), and *μ* that for males, i.e., M/(M+F). If there are *n*_*A*_(*t*) adult females and *n*_*S*_(*t*) female subadults at time *t*, female adults give birth at time *t*+*u*, *u* in [0, 1], will produce,

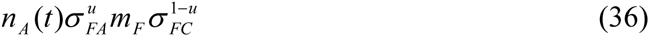

female offspring that survive to time *t*+1 (i.e., a female adult must first survive to time *t*+*u*, then give birth, and then its calf must survive to *t*+1 to be censused at *t*+1; our calculation is an adaptation of Caswell 2001, §6.7.1). Females transitioning from subadulthood to adulthood by giving birth, at time *t*+ *v*, *v* in [0, 1], will produce

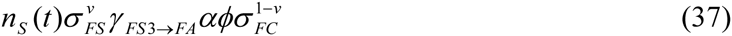

female offspring that survive to time *t*+1. We next took the means of *u* in [0, 1] and *v* in [0, 1] to obtain the per capita fecundities

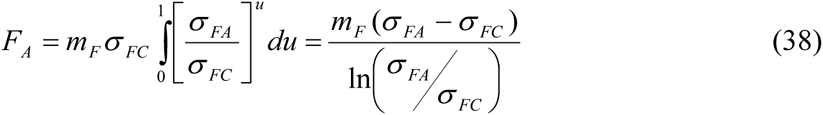

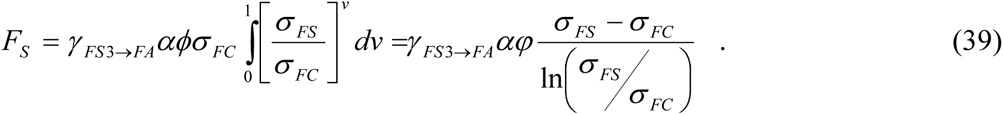

Replacing *m*_*F*_ by *m*_*M*_, *σ*_*FC*_ by *σ*_*MC*_, and *φ* by *μ*, yields the fecundities for male offspring production that give the two nonzero entries for matrix *D*.

The description of our first parametrization (MM1) of the matrix model is now complete.

### MM2 matrix model parametrization

The second parametrization was based on Caswell (2001, §6.1.1). The population projection matrix *A* can be written as the sum *T*+*F*, where the matrix *T* describes transitions and the matrix *F* describes reproduction (Caswell 2001:110). Since each individual’s state is known throughout the study period, one can estimate *T* as follows. For each *t*, one records the number *m*_*ij*_ of individuals in (sub)stage *j* at time *t* that end up in (sub)stage *i* at time *t*+1, where ‘death’ is a possible fate. The matrix *M*_*t*_ = (*m*_*ij*_) contains the matrix *T*_*t*_ as its first *s* rows, where *s* is the number of (sub)stages; its final row contains the mortality information. Caswell (2001, §6.1.1) recommends summing the *M*_*t*_ over *t* to obtain a matrix *M*, and then taking the transition probability *p*_*ij*_ from stage *j* to stage *i* to be the *ij*’th entry of *M* divided by the sum over rows of the entries of *j*’th column of *M* (this estimate is motivated by maximum likelihood estimation). This approach computes *P*’s and *G*’s directly rather than *σ*’s and *γ*’s.

We computed the fecundities using the formulae (38–39) modified so that the transition rates computed in the current method replaced transition and survival rates as computed for the MM1 parametrization. In particular, for *σ*_*FS*_ we used the sum of the transition rates FS3 → FS3 and FS3 → FA, for *σ*_*FC*_ the transition rate FC1a → FC1b, for *σ*_*MC*_ the transition rate MC1a → MC1b, and for *σ*_*FA*_ the transition rate FA → FA. The description of the second parametrization (MM2) is now complete. The actual parametrizations are recorded in Tables 5 and 6.

**Table 5.**
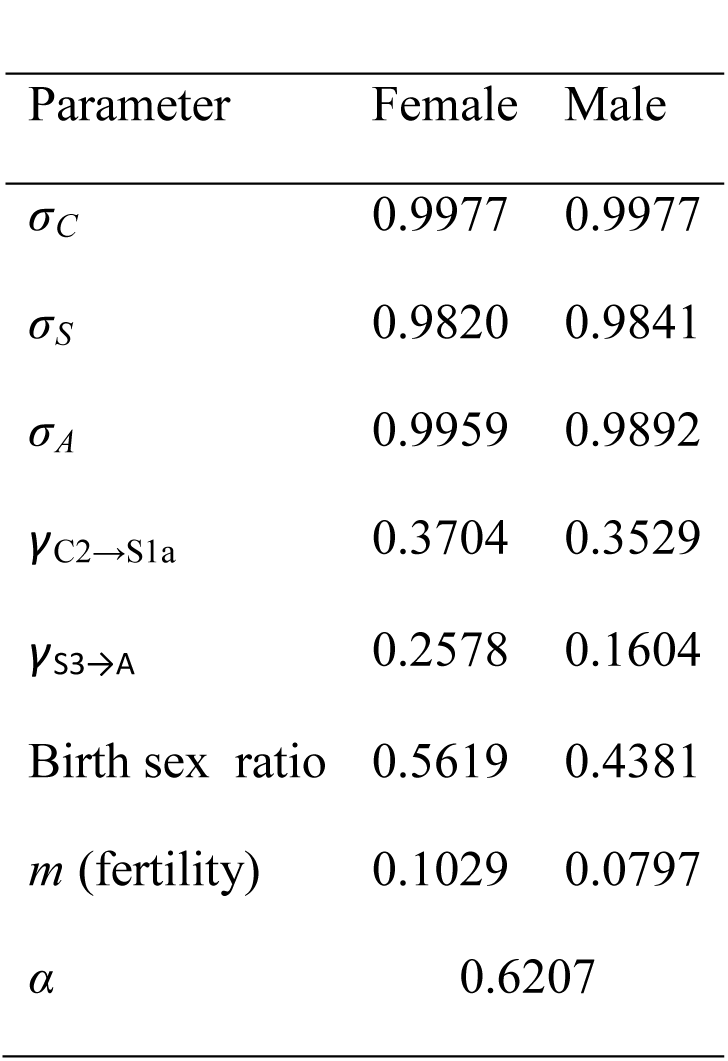
The survival and fertility parameters for the SKKR population. Birth sex ratio is F/(M+F) for females and M/(M+F) for males, where F = number of female births, M = number of male births. The quantity *α* is the probability that a female transitioned from subadulthood to adulthood by giving birth rather than having reached the age of seven years without having given birth (See Table 1).

The most notable difference between MM1 and MM2 is that, for MM2, individuals were more likely to remain within the substage C2 or S3 rather than transition to the next stage, with the consequence that the fecundity *F*_*S*_ was lower for MM2 and individuals reached the adult stage at a slower rate, for both sexes.

**N.B.** Although the matrix models were constructed on a semi-annual time step and with substages for accuracy, our interest is in the biological states C, S, A. Hence, after analyses, substages were collapsed to stages for the purposes of comparison and comparisons were typically made on an annual basis.

**Table 6.** Nonzero entries of the two matrix models MM1 and MM2. The entries in the ‘female’ column yield the female-only part of the model (i.e., the matrix *B* in (29–30)), the entries in the ‘male’ columns for fecundities *F*_S3_ and *F*_A_ are the nonzero entries of the matrix *D* and the remaining entries in the ‘male’ columns given the nonzero entries for the matrix *Q* in (29).

|  | MM1 |  | MM2 |  |
| --- | --- | --- | --- | --- |
| Entry | female | male | female | male |
| $G_{C1a \rightarrow C1b}$ | 0.9977 | 0.9977 | 0.9897 | 0.9899 |
| $G_{C1b \rightarrow C2}$ | 0.9977 | 0.9977 | 1 | 1 |
| $G_{C2 \rightarrow S1a}$ | 0.3666 | 0.3497 | 0.3409 | 0.3173 |
| $G_{S1a \rightarrow S1b}$ | 0.9820 | 0.9841 | 1 | 0.9 |
| $G_{S1b \rightarrow S2a}$ | 0.9820 | 0.9841 | 0.9773 | 1 |
| $G_{S2a \rightarrow S2b}$ | 0.9820 | 0.9841 | 0.9762 | 0.9583 |
| $G_{S2b \rightarrow S3}$ | 0.9820 | 0.9841 | 0.9487 | 1 |
| $G_{S3 \rightarrow A}$ | 0.2549 | 0.1583 | 0.2243 | 0.1087 |
| $P_{C2}$ | 0.6282 | 0.6456 | 0.6591 | 0.6827 |
| $P_{S3}$ | 0.7289 | 0.8262 | 0.7664 | 0.8913 |
| $P_A$ | 0.9959 | 0.9892 | 0.9958 | 0.9878 |
| $F_S$ | 0.0890 | 0.0694 | 0.0775 | 0.0604 |
| $F_A$ | 0.1025 | 0.0795 | 0.1021 | 0.0792 |

### Properties of the two parametrizations

A useful measure of the difference between two population vectors is Keyfitz’s Δ (e.g., Caswell 2001:101). For any two population vectors *X* and *Y*, convert each to a vector of proportions by dividing each component by the sum of that vector’s components. If the resulting vectors of proportions (which sum to one for each vector) are *x* and *y*, then

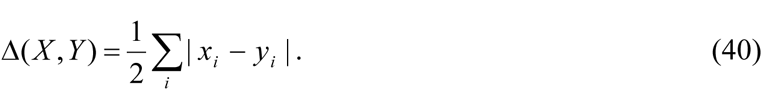

Keyfitz’s Δ has a maximum value of one and is zero when the two vectors coincide.

To compare the two matrix parametrizations, we collapsed the projections of MM1 and MM2 into stage-based population vectors and then computed Keyfitz’s Δ for each parametrization’s projections for each semi-annual census date Jun-99 through Dec-08. Δ = 0.008 in Jun-99, increased monotonically to 0.028 in Dec-00, then decreased monotonically to 0.024 in Dec-03, remained at that value through Dec-05, increased to 0.025 for Jun-06 through Jun-07, and then returned to 0.024 for Dec-07 through Dec-08, with mean 0.0241 and SD 0.0043.Thus, the overall difference between the projections of the two parametrizations is small.

The annual intrinsic rate of increase *r* was 0.1024 (MM1) or 0.0994 (MM2). MM1 and MM2 had the same overall patterns for their 18 eigenvalues and eigenvectors. In order of decreasing magnitude, after the dominant eigenvalue, 1.0525 (MM1) or 1.0510 (MM2), the next two eigenvalues were both real with ratios of 0.99 (MM1 and MM2) and 0.83 (MM1) or 0.89 (MM2) to the dominant. For both parametrizations, the eigenvectors of these two eigenvalues had nonzero components only for the MA, and for the MS3 and MA, components, respectively. The significance of these facts is that the approach to the stable stage distribution (SSD) of the matrix models will be slowest for these two components, i.e., for male adults and MS3s. There follows a complex conjugate pair of eigenvalues, two further distinct real eigenvalues, and two distinct pairs of complex conjugate pairs. Each of these eigenvalues has a single eigenvector. The final eigenvalue (real) has a single linearly independent eigenvector of multiplicity six. The second, third, seventh and last eigenvalues are all real and are the eigenvalues of the matrix *Q*; the fact that the last eigenvalue has algebraic multiplicity six but only geometric multiplicity one reflects the fact that the matrix *Q* is singular. The other eigenvalues are those of the female-only matrix model *B*.

The elasticity of *λ* with respect to adult female survival was the largest elasticity, 0.5396 (MM1) or 0.5349 (MM2), all other elasticities were less than 0.1, and ordered by magnitude in the same way for the two models, with the least being that with respect to the fecundity *F*_*S*_, 0.0059 (MM1) or 0.0055 (MM2). In decreasing order of magnitude, after the highest elasticity comes that with respect to FS3 survival, then that with respect to FC2 survival, then that with respect to each of the transitions FC1a → FC1b, FC1b →FC2, FC2 → FS1a, FS1a → FS1b, FS1b → FS2a, FS2a → FS2b, FS2b → FS3 all of which coincide, then that with respect to the transition FS3 → FA and that with respect to the fecundity *F*_*A*_, which coincide, and finally the smallest, that with respect to the fecundity *F*_*S*_ (see Table 10 for values). The reproductive values of the substages increased from a (normalized) value of 1 for FC1a to a value of 1.81 (MM1) or 1.85 (MM2) for FA (recall that male (sub)stages have zero reproductive value for the Goodman two-sex model). Thus, as expected for a long-lived species, the adult female stage has the greatest influence on demography both as regards the influence of adult female survival on *λ* and reproductive value. As already noted, however, our matrix model does not describe the true asymptotic state of the SKKR population so the elasticity results should not be over interpreted. The stable stage distributions for *stages*, rather than *substages*, are recorded in Table 7, the differences consistent with the previous observation that individuals are slightly more likely to remain as calves or subadults rather than transition to the next stage for MM2 relative to MM1.

**Table 7.** The stable stage distributions (SSD) for MM1 and MM2.

|  | MM1 | MM2 |
| --- | --- | --- |
| stage | SSD | SSD |
| FC | 0.1371 | 0.1395 |
| FS | 0.1550 | 0.1602 |
| FA | 0.2826 | 0.2787 |
| MC | 0.1089 | 0.1122 |
| MS | 0.1413 | 0.1553 |
| MA | 0.1751 | 0.1541 |

Very roughly, given black rhino reproductive behaviour, one expects a typical adult female black rhino to be accompanied by a calf, and to have one prior calf as a SA in the population (for SKKR, mean (± SD) female calf duration was 2.06 ± 0.80 years; mean male calf duration was 2.05 ± 0.83 years; mean female SA duration was 2.7 ± 1.3 years; mean male subadult duration was 3.1 ± 2.0 years). If the birth sex ratio (BSR) is 1:1, then one expects the SSD to be roughly (0.5, 0.5, 1, 0.5, 0.5, x) (assuming the SSD is achieved prior to density dependence sets in), where x < 1 if adult males survive at a lower rate than adult females. Hence, upon normalizing, one gets a SSD of roughly (0.125, 0.125, 0.25, 0.125, 0.125, y), where y ≤ 0.25, except that all the figures (i.e., other than *y*) should be a little larger than stated if x < 1 (*y* < 0.25), and the figure for MS should be a little larger still and that for MA a little smaller as males are subadults longer, on average, than females are, while the proportion for calves should be slightly less to reflect less than perfect reproduction. Large departures from this rough SSD should reflect departures in the BSR from 1:1. For the SKKR population, which produced 48 F to 38 M during 1999 – 2008, the proportions should be a little higher for females than males (comparing calves with calves, SAs with SAs; (48/38) x 0.125 ≈ 0.158 and (38/48) x 0.125 ≈ 0.099). The SSDs derived from the matrix models are consistent with these expectations. Of course, the onset of density dependence would be expected to alter the proportions of stages.

For each parametrization, the matrix model projections approached their SSDs over the period Jun-99 through Dec-08, with Keyfitz’s Δ between the projection and the SSD strictly decreasing from 0.227 (MM1) or 0.208 (MM2) to 0.001 (MM1) or 0.004 (MM2), respectively. Thus, the transient behaviour present in the matrix model projections during the period of interest basically consisted of the ‘smooth’ extinguishing of the deviation from SSD present in the initial population vector (i.e., SKKR in Dec-98). In particular, for the MM1 parametrization, the damping ratios (Caswell 2001, §4.7.1) for the second and third largest eigenvalues were 1.06 and 1.27, respectively, so that exponential damping of their eigenvectors had half-lives of 11.2 and 2.9 years, respectively. Thus, though the proportion of adult males is somewhat slow to approach the SSD proportion (the eigenvector of the second largest eigenvalue has male adults as its only nonzero component), all other eigenvectors are damped fairly rapidly, implying that all other (sub)stages approach their SSD proportions fairly rapidly. For MM2, the half-life for the second largest eigenvalue is the same as for MM1 but that for the third largest eigenvalue is 4.2, slightly larger, implying that MS3 approaches its SSD proportion a little slower than for MM1, as noted previously. All these half-lives, however, are just a fraction of black rhinoceros lifespan.

### Matrix model analyses and results

Our primary interest in the matrix model was in modelling the dynamics of the stage-structured population after introductions ceased, from Dec-98 through Dec-08. The SKKR population was still a young population. In particular, no rhinoceros born in GFFR died of old age during the study. Thus, though matrix entries of MM1 and MM2 might be plausibly considered representative of the dynamics during the study, as adults age, and deaths due to old age become common, the survival rate of adults will decrease below that during the study period. Thus, the asymptotic dynamics of the matrix model should not be confused with the asymptotic dynamics of the actual population, even in the absence of density dependence in the actual population. Thus, the asymptotic dynamics of the two parametrizations are only indicative of the model and of how the population might have been expected to behave in the long term had nothing else changed, which as noted is unrealistic. The asymptotic properties of the matrix model then are of interest as indicators of how well the matrix model describes an exponentially growing population of large herbivores (‘slow’ mammals) with adult survival rates that are somewhat too high in the long run (e.g., Brodie *et al*. 2011 estimated adult female survival as 0.944, 95% CI = 0.920 – 0.962 and adult male survival as somewhat lower but larger than 0.9). We did not use the matrix model to forecast population dynamics beyond the study period.

The five rhinoceros removed from SKKR in May-06 belonged, as of Jun-06, to the following substages: one to each of FS2a, FS2b, MS3, and two to FS3. The corresponding substage population vector was projected forward from Jun-06 by each of MM1 and MM2 and, for Dec-06, Dec-07, and Dec-08, the projected quantities rounded to integers so as to obtain biologically sensible projections (moreover, since the exported male did not turn eight years old until 2009 it was retained as a subadult throughout), resulting in the following stage-based population vectors (FC, FS, FA, MC, MS, MA):

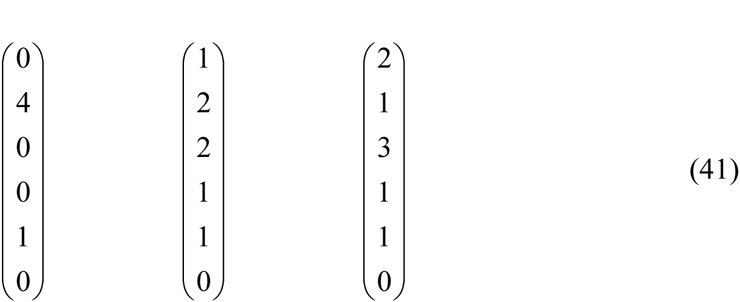

for Dec-06, Dec-07, and Dec-08, respectively, for both MM1 and MM2. The final vector for Dec-08 was slightly ambiguous for the MM2 projection, as an alternative interpretation was that the population vector for Dec-08 was the same as that for Dec-07, but we chose the option in (41) as a consensus result and in order to have a parametrization-independent result. These three vectors were added to the observed SKKR population vectors for Dec-06, Dec-07, and Dec-08. The resulting augmented population vectors, together with the observed SKKR population vectors for each December from 1998 through 2005, will henceforth be referred to as the SKKR population vectors and were the population vectors to which the matrix projections were compared. Thus, the actual SKKR substage population vector for Dec-98 was projected semi-annually by each of MM1 and MM2 up to Dec-08. For each December, 1999 through 2008, these projections were collapsed to stage-based population vectors and compared to the actual stage-based population vectors. In addition the projected number of additions each year (i.e., born during that year and survived to the end of that year), by sex, were recorded and compared to the actual number of additions, by sex, per year. In addition to direct comparison, we computed Keyfitz’s Δ. The results for both parametrizations are presented in Table 8. Figure 5 shows plots of Keyfitz’s Δ.

**Figure 5.**
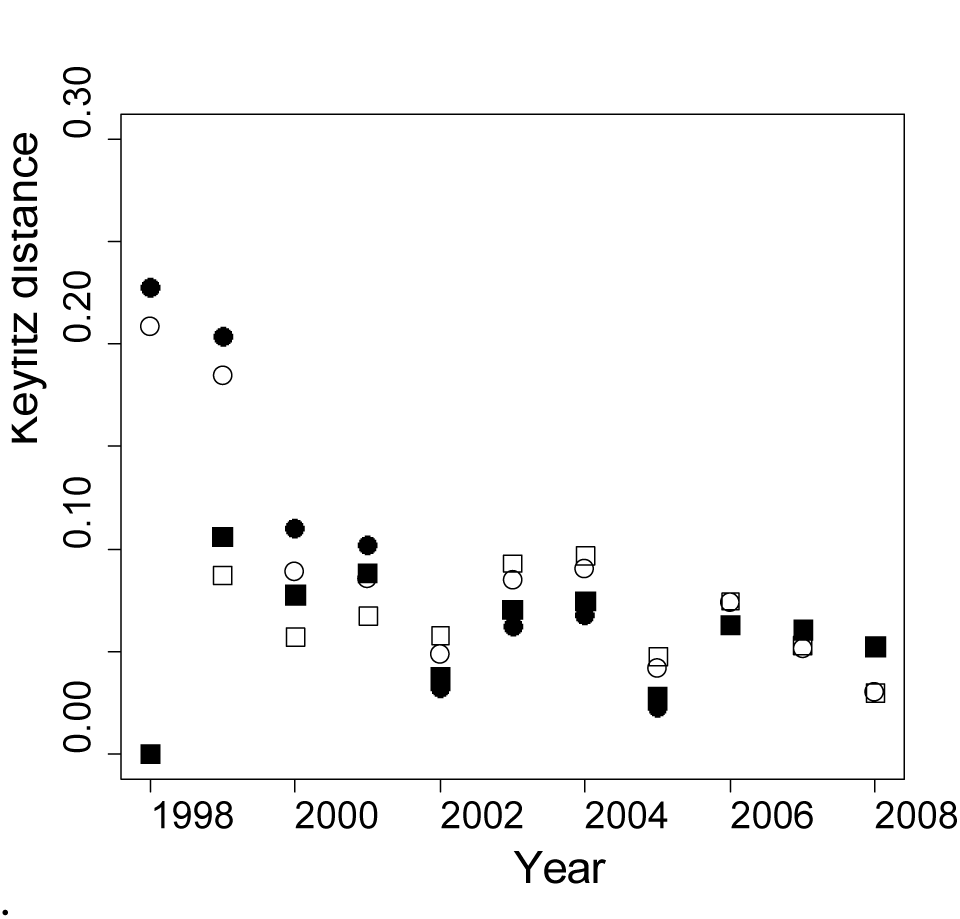
Keyfitz’s Δ for the SKKR population vector and: the SSD of MM1 (solid circles); the SSD for MM2 (open crcles); the projections of MM1 (solid squares); the projections MM2 (open squares). Unlike the projections’ monotone convergence on their SSDs (described on p. 34), the SKKR population vectors, though eventually close to SSDs and model projections, exhibited mild fluctuations relative to them.

**Table 8.** The second and fourth columns list the matrix model projections for the end of year for parametrizations MM1 and MM2, respectively; the third column lists the SKKR population vectors (augmented by the exports of 2006 plus their projections for 2007 and 2008) at the end of year; all these population vectors are formatted as the transpose of the row vector (FC, FS, FA, MC, MS, MA); column six lists the number of female and male recruits each year (augmented by the projected additions from the 2006 exports) as a column vector 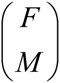; the fifth and seventh columns list the number of projected female and male recruits *M* each year (recruits are offspring of the year surviving to the end of the year), in the same format as column six, for MM1 and MM2, respectively; the final column lists Keyfitz’s Δ for the SKKR population vector of column three and the MM1 Projection in column two (Δ_1_), and the MM2 projection in column four (Δ_2_) (also plotted in Fig. 5).

| Year | MM1 | SKKR | MM2 | MM1 | SKKR | MM2 | $\Delta_1$ |
| --- | --- | --- | --- | --- | --- | --- | --- |
| | Pop Vec | Pop Vec | Pop Vec | recruits | recruits | births | $\Delta_2$ |
| 1999 | 5.6 | 6 | 5.6 | 3.1 | 4 | 2.9 | 0.1059 |
|  | 9.4 | 13 | 9.6 | 2.4 | 0 | 2.2 | 0.0870 |
|  | 11.6 | 9 | 11.2 |  |  |  |  |
|  | 5.0 | 3 | 4.9 |  |  |  |  |
|  | 9.4 | 10 | 9.9 |  |  |  |  |
|  | 5.5 | 5 | 5.0 |  |  |  |  |
| 2000 | 7.2 | 7 | 7.0 | 3.6 | 1 | 3.4 | 0.0774 |
|  | 8.0 | 9 | 8.5 | 2.8 | 0 | 2.7 | 0.0570 |
|  | 14.6 | 12 | 14.0 |  |  |  |  |
|  | 5.9 | 3 | 5.8 |  |  |  |  |
|  | 9.1 | 9 | 9.9 |  |  |  |  |
|  | 7.3 | 6 | 6.4 |  |  |  |  |
| 2001 | $\begin{pmatrix} 8.1 \\ 8.5 \\ 16.4 \\ 6.5 \\ 9.1 \\ 9.1 \end{pmatrix}$ | $\begin{pmatrix} 6 \\ 9 \\ 17 \\ 7 \\ 12 \\ 6 \end{pmatrix}$ | $\begin{pmatrix} 8.0 \\ 8.8 \\ 15.8 \\ 6.5 \\ 10.1 \\ 7.9 \end{pmatrix}$ | $\begin{pmatrix} 3.7 \\ 2.9 \end{pmatrix}$ | $\begin{pmatrix} 4 \\ 7 \end{pmatrix}$ | $\begin{pmatrix} 3.6 \\ 2.8 \end{pmatrix}$ | 0.0879 |
|  |  |  |  |  |  |  | 0.0675 |
| 2002 | $\begin{pmatrix} 8.9 \\ 9.6 \\ 18.2 \\ 7.1 \\ 9.8 \\ 10.5 \end{pmatrix}$ | $\begin{pmatrix} 7 \\ 9 \\ 18 \\ 7 \\ 8 \\ 10 \end{pmatrix}$ | $\begin{pmatrix} 8.9 \\ 9.7 \\ 17.6 \\ 7.2 \\ 10.8 \\ 9.1 \end{pmatrix}$ | $\begin{pmatrix} 4.1 \\ 3.2 \end{pmatrix}$ | $\begin{pmatrix} 3 \\ 0 \end{pmatrix}$ | $\begin{pmatrix} 4.0 \\ 3.1 \end{pmatrix}$ | 0.0372 |
|  |  |  |  |  |  |  | 0.0579 |
| 2003 | $\begin{pmatrix} 9.7 \\ 10.9 \\ 20.0 \\ 7.8 \\ 10.6 \\ 11.9 \end{pmatrix}$ | $\begin{pmatrix} 12 \\ 10 \\ 20 \\ 7 \\ 7 \\ 14 \end{pmatrix}$ | $\begin{pmatrix} 9.7 \\ 10.9 \\ 19.3 \\ 7.8 \\ 11.7 \\ 10.4 \end{pmatrix}$ | $\begin{pmatrix} 4.5 \\ 3.5 \end{pmatrix}$ | $\begin{pmatrix} 7 \\ 4 \end{pmatrix}$ | $\begin{pmatrix} 4.3 \\ 3.3 \end{pmatrix}$ | 0.0703 |
|  |  |  |  |  |  |  | 0.0929 |
| 2004 | $\begin{pmatrix} 10.7 \\ 12.2 \\ 22.0 \\ 8.5 \\ 11.6 \\ 13.4 \end{pmatrix}$ | $\begin{pmatrix} 13 \\ 11 \\ 23 \\ 7 \\ 8 \\ 15 \end{pmatrix}$ | $\begin{pmatrix} 10.7 \\ 12.2 \\ 21.2 \\ 8.6 \\ 12.7 \\ 11.7 \end{pmatrix}$ | $\begin{pmatrix} 5.0 \\ 3.9 \end{pmatrix}$ | $\begin{pmatrix} 5 \\ 3 \end{pmatrix}$ | $\begin{pmatrix} 4.7 \\ 3.7 \end{pmatrix}$ | 0.0743 |
|  |  |  |  |  |  |  | 0.0967 |
| 2005 | $\begin{pmatrix} 11.8 \\ 13.5 \\ 24.4 \\ 9.4 \\ 12.7 \\ 15.0 \end{pmatrix}$ | $\begin{pmatrix} 12 \\ 13 \\ 24 \\ 10 \\ 10 \\ 15 \end{pmatrix}$ | $\begin{pmatrix} 11.7 \\ 13.5 \\ 23.4 \\ 9.4 \\ 13.8 \\ 13.0 \end{pmatrix}$ | $\begin{pmatrix} 5.5 \\ 4.3 \end{pmatrix}$ | $\begin{pmatrix} 4 \\ 6 \end{pmatrix}$ | $\begin{pmatrix} 5.2 \\ 4.1 \end{pmatrix}$ | 0.0276 |
|  |  |  |  |  |  |  | 0.0475 |
| 2006 | 13.1 | 12 | 13.0 | 6.1 | 8 | 5.8 | 0.0628 |
|  | 14.9 | 18 | 14.9 | 4.8 | 7 | 4.5 | 0.0744 |
|  | 27.1 | 27 | 25.9 |  |  |  |  |
|  | 10.4 | 14 | 10.4 |  |  |  |  |
|  | 13.8 | 10 | 15.0 |  |  |  |  |
|  | 16.7 | 17 | 14.5 |  |  |  |  |
| 2007 | 14.6 | 12 | 14.3 | 6.8 | 3 | 6.4 | 0.0604 |
|  | 16.5 | 18 | 16.5 | 5.3 | 6 | 5.0 | 0.0532 |
|  | 30.0 | 29 | 28.6 |  |  |  |  |
|  | 11.6 | 16 | 11.5 |  |  |  |  |
|  | 15.2 | 14 | 16.4 |  |  |  |  |
|  | 18.6 | 16 | 16.1 |  |  |  |  |
| 2008 | 16.2 | 16 | 15.8 | 7.5 | 9 | 7.1 | 0.0522 |
|  | 18.2 | 20 | 18.2 | 5.9 | 5 | 5.5 | 0.0296 |
|  | 33.3 | 32 | 31.6 |  |  |  |  |
|  | 12.8 | 14 | 12.7 |  |  |  |  |
|  | 16.8 | 20 | 18.0 |  |  |  |  |
|  | 20.7 | 16 | 17.9 |  |  |  |  |

We offer the following account of the results recorded in Table 8. In this account, stages such as FC will refer to the SKKR population vector in comparison to the projections. ‘Recruits’ are offspring of the year that survive to the end of the year; for the SKKR population, probability of death in the first year after birth was negligible, so recruits are essentially births. First note that while transitions from MS3 to MA occurred in SKKR by a male reaching age eight (Table 1), in the model they occurred by probability and the matrix model does not capture the actual distribution of male ages in stage S3 of the SKKR population. In 1999, FS was high and FA low compared to the projection, so FS transitioned to FA less than expected by the model while MC was low due to absence of male recruits in 1999; this state of affairs persisted for 2000. The low number of recruits in 2000 combined with the high number of males born in 2001 resulted in FC lagging behind projections but MC having caught up; FS to FA transitions resulted in rough agreement for these stages but MS grew larger than projected, resulting in a deficit for MA. During 2002, MS to MA transitions brought closer agreement with projections and the remaining stages maintained their status. The high number of female recruits in 2003 put FC ahead of projections but MS began to lag behind projections, reflecting the absence of male recruits in 1999–2000. MA moved ahead of projections indicating a higher transition rate of MS to MA than projected in 2003. In 2004, little had changed except that male recruits are slightly less than projected. In 2005 there was close agreement between projections and SKKR, except that MS still lagged. The discrepancy in FC from 2003 – 2004 has been eliminated. A high number of recruits occurred in 2006 compared to projections; by the end of that year FC lagged by one behind projection, and FS moved ahead of projections (a higher rate of FC to FS transitions than projected, induced by the higher number of actual recruits therefore causing more calves to become independent) but MC was ahead of projections while MS still lagged. The lag in MS 2002 – 2006 appears to reflect the absence of male recruits in 1999 – 2000, the fact that MS deaths, although few, were concentrated in 2003 – 2005 (whereas FS deaths were more spread out over time), and the higher rate of MS to MA transitions noted for 2003. In 2007, FC lagged further behind projection due to the lower number of female recruits than projected, FS was more in line with projections, and the state of affairs for the other stages was basically unchanged. In 2008, a higher number of female recruits than projected occurred, and FC was now in line with projections, while FS had slightly increased its advance over projection since 2007, the excess deriving from the greater-than-projected number of female recruits in 2006. The higher number of male recruits than projected for 2005 – 2007 maintained MC ahead of projections and also pushed MS ahead of projections, but these male recruits had not yet affected MA.

In summary, we propose that the differences in number and sex between the actual annual recruits and the matrix-model projected recruits, and the resulting knock-on effects as calves transition to subadults and subadults to adults explain much of the discrepancy between the SKKR population vectors and matrix-model projections. The fluctuations in Keyfitz’s Δ, both between the SKKR population vectors and the projections and between the former and the SSDs (Fig. 5), are greater than the transient behaviour manifest in the matrix models themselves for the period 98 – 08 (p. 34). Nevertheless, as measured by Keyfitz’s Δ, the differences between the actual SKKR population vectors and the matrix model projections is small, less than 0.1 after 1999.

### Transient dynamics

We noted that the asymptotic properties of the matrix model are of limited interest as the SKKR population was still young and adult mortality rates will increase after 2008 due to individuals dying of old age and eventually density dependence will have some effect also. The primary interest then of the asymptotic properties of the matrix model was in assessing closeness of the matrix model projections during 98–08 to the SSD as an indication of transient dynamics, i.e., deviations from the SSD due to the initial population vector. Koons *et al*. (2005) drew attention to the fact that sensitivities of transient growth may differ from sensitivities of asymptotic growth. The period Dec-98 through Dec-08 is of interest not only for understanding transient dynamics in their own right but also because during this period removals commenced to source reintroductions elsewhere, so any transient behaviour may have consequences for such harvesting.

We used the matrix model (we conducted the following analyses for each parametrization MM1 and MM2; results were very similar and we only report those for MM1) to project the SKKR substage population vector for each December, 1998 through 2007, through two time steps to the following December, compared that projection with the SKKR population vector for the December to which the projection was made using Keyfitz’s Δ applied to the stage-based population vectors (i.e., after collapsing substages), and computed the transient annual growth rate for each such projection as

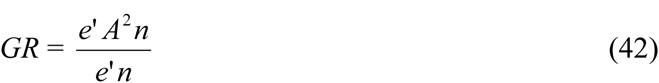

where *n* is the SKKR substage population vector, *e* is a column vector all of whose components are 1, and *e*’ denotes its transpose. Note that for this computation, the SKKR vectors did not need to be augmented with the exports and their projections to 2007 – 2008, except that the exports themselves were retained for the SKKR 2006 population vector that was compared to the projection from the 2005 SKKR population vector. The exports were excluded from the 2006 SKKR population vector that was projected to 2007. For the 10 projections, the mean *GR* ± SD was 1.1111 ± 0.0084, and the range from 1.0967 (2006) to 1.1267 (2000), as compared to the asymptotic *annual* growth rate (*λ*) of 1.1078 of MM1, (Table 9). These results are consistent with the fact (Fig. 5) that SKKR population vectors during this period did not stray much from the SSD.

**Table 9.** Transient annual growth rates computed from (42) from actual SKKR substage population vectors, each December, 1998–2007.

| Year | Growth rate |
| --- | --- |
| 1998 | 1.1078 |
| 1999 | 1.1191 |
| 2000 | 1.1267 |
| 2001 | 1.1127 |
| 2002 | 1.1140 |
| 2003 | 1.1166 |
| 2004 | 1.1118 |
| 2005 | 1.1030 |
| 2006 | 1.0967 |
| 2007 | 1.1026 |

Values of *GR* below *λ* could sound a warning to managers planning to remove animals during the coming year that the population vector is currently expected to perform below the stable rate *λ*. For SKKR, that condition pertained from 2005–2007, with values smaller by -0.4, -1.0, and -0.5% of the value of *λ*, respectively. The smallest value for *GR* occurred for the projection from Dec-06 to Dec-07, after the removals of the five SAs. These five were expected to have contributed two new animals to the population during that year according to equation (40), which might account for that lowest anticipated annual recruitment from 2006 to 2007.

Keyfitz’s Δ between the projection to a given December and that December’s SKKR population vector (after collapsing substages) averaged 0.054, with a SD of 0.025 and a low of 0.020 for the projection from 2003 to 2004 (Fig. 6). These values allow a retrospective assessment of actual population performance versus anticipated performance based on the matrix model. For example, note that the largest value of *GR* occurred for the projection from Dec-00 to Dec-01, which is also the projection for which the disparity between actual and predicted performance is greatest as measured by Δ. During that year, there were 11 animals recruited to the population but the projected number was only about three, so the SKKR population actually outperformed the annual projection that year. The lowest value of Δ occurred for the projection Dec-03 to Dec-04, for which *GR* had its third highest value; for this year actual additions (eight) and anticipated additions coincided closely. For the three years in which *GR* was less than *λ*, Δ was never large than 0.05 so actual population performance was similar to predictions. Further years of data would have been interesting to see if there was a signal of a trend in these values of *GR* less than *λ*.

**Figure 6.**
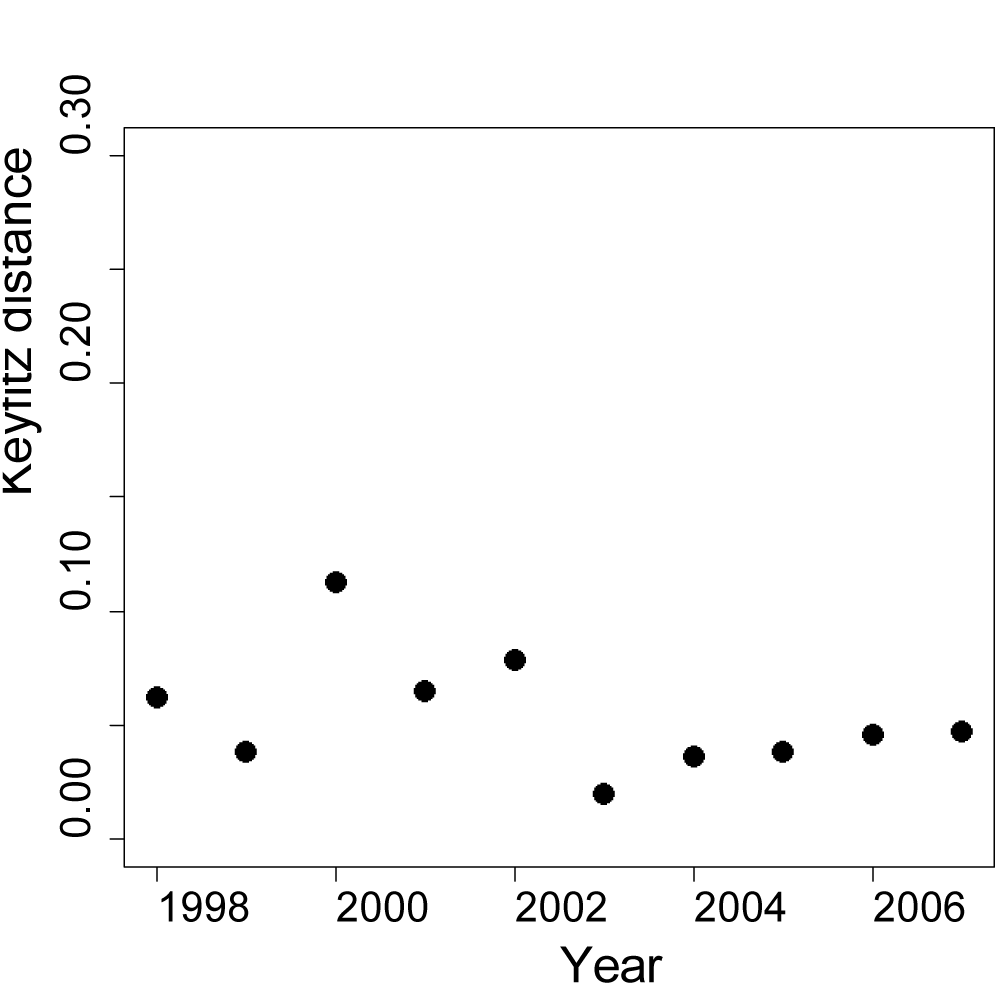
Plot of Keyfitz’s Δ between the projections over one year (two time steps) of matrix model MM1 of SKKR population vectors for each December, 1998–2007, and the actual SKKR population vector the following December.

**Figure 7.**
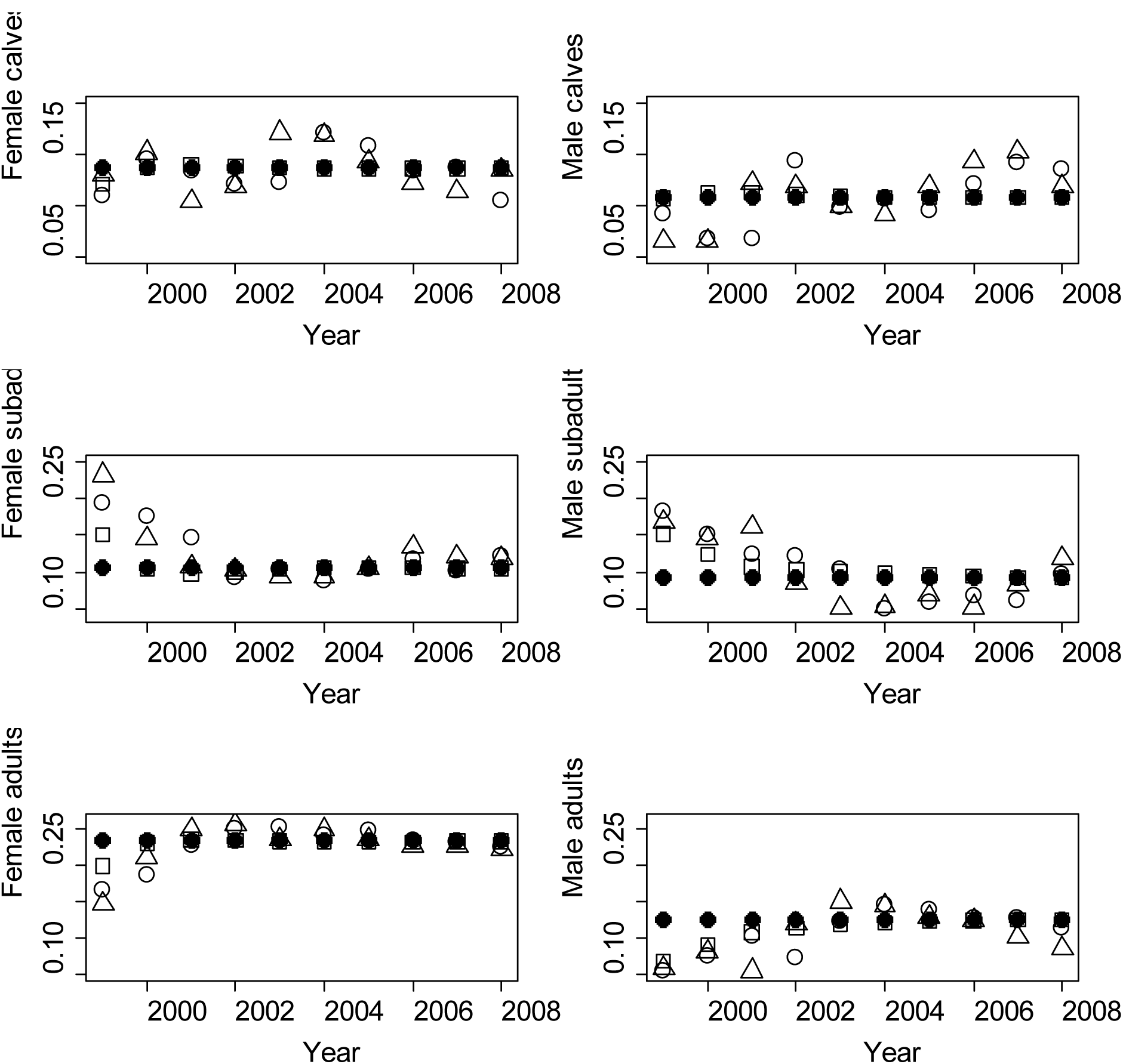
The proportion of each stage-and-sex, is plotted against year, Dec-99 through Dec-08 for: the matrix model (MM1) stable stage distribution (•) the SKKR population vector (Δ); the matrix model (MM1) projections of the SKKR population vector in Dec-98 (□); and the matrix model projection (MM1) of the SKKR population vector from the previous December to that year’s December (○). The SKKR population vector in Dec-98 was (FC,FS,FA,MC,MS,MA) = (5,11,8,3,11,4) with proportions (0.12,0.26,0.19,0.07,0.26,0.10) versus the SSD of (0.14,0.155,0.28,0.11,0.14,0.175).

In Fig. 7, the SKKR population vectors exhibit the least convergence on MM1’s SSD, MM1’s projections from Dec-98 the most, with the annual projections reflecting the proportions of the SKKR vector of the previous year. As argued on pp. 39–40, it is the discrepancies in actual recruits from projected recruits, manifest in the fluctuations of the proportions of calf stages that result in the deviation of the actual population dynamics from those of the model.

Koons *et al*. (2005) computed the sensitivities to entries *a*_*ij*_ of *A* of the growth over a single time step at arbitrary time *t*-1

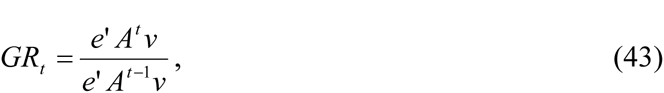

where *v* is the population vector at time *t*-1, by calculating the partial derivatives of *GR*_*t*_ to *a*_*ij*_. As an aside, we note that the resulting sensitivities to *a*_*ij*_ of growth *GR*_1_ over one time step is just the proportion of the *j*’th stage in the initial population vector, independently of the form of *A* or the size of the time step, indicating the importance of the initial population vector. If the initial population vector is the SSD, the growth is λ and one might expect to obtain the sensitivities of the asymptotic growth rate λ, but the two formulae only agree if the reproductive value vector *w* of *A* equals *e*, which one would not expect. The difference stems from the fact that in computing the sensitivities of the asymptotic growth rate, the population vector *v* is the SSD and thus depends on *a*_*ij*_ too, whereas for the transient growth rate *v* is fixed. Moreover, if *v*_*j*_ = 0, then the transient sensitivity with respect *a*_*ij*_, any *i*, is zero, because no variation in *a*_*ij*_ can effect *GR*_1_ when *v*_*j*_ is zero.

We adapted Koons *et al*.’s notion of sensitivities of transient growth to our purposes by taking the partial derivatives of (42) with respect to entries of *A*, which results in the following formula:

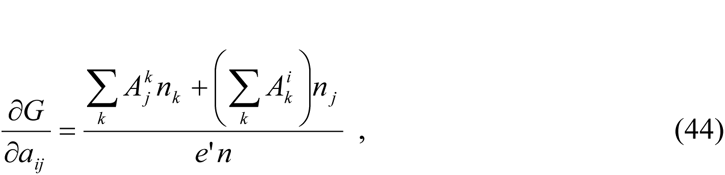

where here 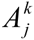 denotes the entry of the matrix A in the *j*’th row and *k*’th column and *n*_*k*_ denotes the *k*’th component of *n*. These sensitivities indicate the dependence on the matrix entries of the transient one-year growth projections from actual SKKR states, which might warn the manager of unusual transient demographics. We converted sensitivities to elasticities in the usual manner (Caswell 2001:226). Note that our *GR* in (42) is homogeneous of degree two in the entries of *A*, so by Euler’s formula (Caswell 2001:229), our elasticities will sum to two.

**Table 10.** Means ± SD for the sensitivities and elasticities of *GR* (42) computed by (44) and sensitivities and elasticities of the dominant eigenvalue of MM1.

| Matrix entry | Sensitivity of $\lambda$ | Mean (sensitivity of GR) ± SD | Elasticity of $\lambda$ | Mean (elasticity of GR) ± SD |
| --- | --- | --- | --- | --- |
| $F_S$ fecundity | 0.070 | $0.178 \pm 0.117$ | 0.0059 | $0.0142 \pm 0.0086$ |
| $F_A$ fecundity | 0.315 | $0.558 \pm 0.067$ | 0.0307 | $0.0515 \pm 0.0062$ |
| FC1a → FC1b | 0.039 | $0.062 \pm 0.019$ | 0.0366 | $0.0552 \pm 0.0169$ |
| FC1b → FC2 | 0.039 | $0.057 \pm 0.028$ | 0.0366 | $0.0512 \pm 0.0247$ |
| FC2 → FC2 | 0.091 | $0.157 \pm 0.040$ | 0.0542 | $0.0888 \pm 0.0222$ |
| FC2 → FS1a | 0.105 | $0.156 \pm 0.040$ | 0.0366 | $0.0515 \pm 0.0129$ |
| FS1a → FS1b | 0.039 | $0.052 \pm 0.018$ | 0.0366 | $0.0460 \pm 0.0160$ |
| FS1b → FS2a | 0.039 | $0.043 \pm 0.029$ | 0.0366 | $0.0386 \pm 0.0259$ |
| FS2a → FS2b | 0.039 | $0.042 \pm 0.022$ | 0.0366 | $0.0369 \pm 0.0199$ |
| FS2b → FS3 | 0.039 | $0.051 \pm 0.039$ | 0.0366 | $0.0453 \pm 0.0344$ |
| FS3 → FS3 | 0.119 | $0.191 \pm 0.116$ | 0.0825 | $0.1249 \pm 0.0344$ |
| FS3 → FA | 0.127 | $0.194 \pm 0.118$ | 0.0307 | $0.0444 \pm 0.0268$ |
| FA → FA | 0.570 | $0.606 \pm 0.074$ | 0.5396 | $0.5436 \pm 0.0671$ |
| $M_S$ fecundity | 0 | $0.178 \pm 0.108$ | 0 | $0.0111 \pm 0.0067$ |
| $M_A$ fecundity | 0 | $0.558 \pm 0.067$ | 0 | $0.0399 \pm 0.0048$ |
| MC1a → MC1b | 0 | $0.054 \pm 0.019$ | 0 | $0.0485 \pm 0.0172$ |
| MC1b → MC2 | 0 | $0.049 \pm 0.038$ | 0 | $0.0438 \pm 0.0342$ |
| MC2 → MC2 | 0 | $0.116 \pm 0.050$ | 0 | $0.0672 \pm 0.0291$ |
| MC2 → MS1a | 0 | $0.115 \pm 0.049$ | 0 | $0.0362 \pm 0.0156$ |
| MS1a → MS1b | 0 | $0.036 \pm 0.014$ | 0 | $0.0317 \pm 0.0124$ |
| MS1b → MS2a | 0 | $0.029 \pm 0.023$ | 0 | $0.0254 \pm 0.0201$ |
| MS2a → MS2b | 0 | $0.027 \pm 0.015$ | 0 | $0.0239 \pm 0.0137$ |
| MS2b → MS3 | 0 | $0.033 \pm 0.029$ | 0 | $0.0295 \pm 0.0260$ |
| MS3 → MS3 | 0 | $0.190 \pm 0.094$ | 0 | $0.1411 \pm 0.0688$ |
| MS3 → MA | 0 | $0.191 \pm 0.094$ | 0 | $0.0271 \pm 0.0132$ |
| MA → MA | 0 | $0.317 \pm 0.068$ | 0 | $0.2827 \pm 0.0609$ |

For most nonzero entries of *A*, the sensitivities varied without obvious pattern across the years, except that, for both sexes, those for subadult fecundity (*F*_*S*_ and *M*_*S*_) and the transition probabilities from S to A (*G*_FS3→FA_ and *G*_MS3→MA_) tended to decrease, while those for female adult fecundity (*F*_*A*_ and *M*_*A*_) and adult survival (*P*_*FA*_ and *P*_*MA*_) tended to increase, as one might expect for a long-lived species. The same patterns were observed for the elasticities. One can think of a given annual projection of an SKKR population vector and the corresponding *GR* as the growth expected over the next year. Any unusual departure in the rankings of sensitivities and elasticities of *GR* to the matrix entries of the female component of the matrix model to those of *λ* could serve as a warning of unusual transient dynamics, which might be relevant to management practices. In the present case, there do not appear to be any warning bells of very unusual demography. The patterns in sensitivities and elasticities were similar for both transient and asymptotic growth rates.

In summary, the matrix model projections converged towards the SSD over the modelled period 1998–2008, indicating the dynamics were only mildly transient during this period in the sense that the projections were not already in the SSD. Moreover, the projected annual growth rates (42) each year 1998–2007 differed by less than 1.8% from the asymptotic growth rate *λ* (Table 7). The transient sensitivities and elasticities (Table 10) did not indicate any surprising departures from expectations based on asymptotic dynamics.

## 4. Demographic Stochasticity of the Structured Population Dynamics

Sæther *et al*. (1998b) used a fairly complicated procedure (some details of which were unpublished) to estimate demographic and environmental stochasticity for brown bears accounting for their population structure. Engen *et al*. (2005) developed a simpler method based on matrix models and obtained, for long-lived vertebrates that produce only a single offspring per breeding occasion, and assuming no relationship between reproduction and subsequent adult survival and no environmental stochasticity, an equation that estimates demographic stochasticity from the deterministic (female-only) matrix model presumed to underlie the dynamics. It appears plausible to apply this equation to the female segment of our matrix model. The demographic assumptions apply to black rhinoceros. While an absence of environmental stochasticity will not be generally valid for black rhinoceros, the estimate of environmental stochasticity for SKKR using the scalar model of exponential growth in section 2 indicates it was negligible, a result we shall argue in the Discussion is consistent with our previous studies of SKKR. Moreover, the previous estimate of demographic stochasticity reflected variation in fecundity (at least 86%) more than survival so restricting to the female-only segment of the population focuses on the most important source of demographic stochasticity. The method of Engen *et al*. (2005) will provide a more refined estimate of demographic stochasticity than the estimate in section 2 by accounting for differences between individuals due to stage structure.

Engen *et al*.’s equation (13) is written down for an age-structured population but it is a simple matter to extend it to a stage-based matrix model. Recall that λ is the same for the full and female-only matrix models. Let (*u*_*i*_) be the SSD of a female-only matrix model and let (*v*_*i*_) be the reproductive value vector, normalized to have unit scalar product with the SSD. Then,

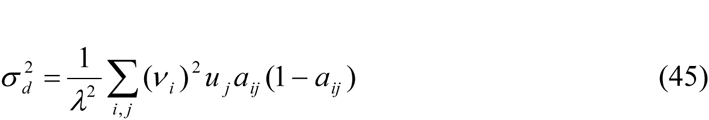

where the summation is over nonzero entries of the projection matrix *A* = (*a*_*ij*_). Since our matrix model was constructed with a semi-annual time step, we applied (45) to *Z* = *A*^2^, which is the projection over one year.

The value obtained for matrix model MM1 was 0.105 and 0.106 for MM2 for semi-annual time steps, so we only report the result for *A*^2^ for MM1, which was 0.204.

## DISCUSSION

Consistent with Nelder (1961), estimates of *K* and especially *θ* were poor, and often useless, when fitting the time series of abundances to the various versions of the generalized logistic. Exponential growth can be mimicked by the generalized logistic with large values of *K* and/or *θ*, resulting in a redundancy in these parameters when fit to exponential-growth-like data. Fitting exponential growth data to a generalized logistic will force the inflexion point (5) of the latter to be located beyond the range of the actual data. The estimates of *K* and their CVs obtained from the models CGL and SGL with fixed values of *θ* were increasingly more precise as *θ* increased, i.e., as these models become more threshold-like, reflecting the fact that increasingly threshold-like models can be fit to exponential growth with an inflexion increasingly just beyond the final census value. Thus, the competitiveness of these models, as measured by AIC_c_, merely reflected the degree to which they represent exponential growth prior to their inflexion point. The generalized logistic will be unreliable for modelling population dynamics when the data samples only abundances near *K* (Polansky *et al*. 2009; Clark *et al*. 2010) or only abundances prior to the onset of density dependence (our study) but does appear to be useful for data across the range of population growth (Eberhardt *et al*. 2008).

Semi-annual census data over 22.5 years for the reintroduced and expanding black rhinoceros SKKR population was unambiguously best fit, amongst scalar population models based on the generalized logistic (Table 2), by the discrete-time exponential model with multiplicative error. As likelihood models of this data, however, this model is indistinguishable from the model of Dennis *et al*. (1991) and of the continuous-time stochastic exponential growth model (Levins 1969, Tuckwell 1974). Thus our results do not discriminate between continuous-and discrete-time versions of stochastic exponential growth but do favour multiplicative over additive error. One expects process noise to be multiplicative on population growth if it is additive on vital rates (Turchin 2003:184). The non-competitive performance of the naïve continuous-time exponential growth model indicated that process noise contributed to the dynamics.

The naïve continuous-time models were favoured over the discrete-time exponential model for the annual (December) census data, however. Residuals of the fit of models indicated that differences between the results for the two time steps was an artifact of the annual time step rather than biologically informative (Figs. 3–4). It appears that stochasticity contained in the semi-annual censuses tended to average out over the annual interval. Hence, censuses limited to longer intervals can misrepresent the dynamics of populations with asynchronized reproduction and no natural time step.

The absence of information in our data regarding *K* is consistent with the threshold model (6) adopted in Emslie (2001), but does not confirm it. Whether it is more likely that the dynamics of SKKR conform to the threshold model rather than a generalized logistic model, with a value of *θ* exhibiting a slower decline in pgr after the inflexion, depends on how close the SKKR population approached its carrying capacity by December 2008. The highest (local) density for black rhinoceros reported by Owen-Smith (1988) was 1.6/km^2^. Employing this figure as a mean density for SKKR yields 352 as an upper bound for *K*. Ignoring stochasticity, it would take 11.5 years to reach this figure from the final census of 110 via exponential growth with the value of *r* obtained from our best model. Stochasticity and removals aside, then, one could expect to discern the form of density dependence for SKKR within another 11.5 years, either a relatively sudden cessation in exponential growth for a threshold model or a more gradual decline in pgr consistent with the generalized logistic with a value of *θ* larger than one, but smaller than about ten.

For annual survey data for two black-rhinoceros populations that exhibited leveling off of population size, Cromsigt *et al*. (2002) employed discrete-time models with additive error (interpreted as observation error) and obtained good fits to their data with the DGL form with estimates for *θ* of 10 and 28 (no SEs reported). For three black-rhinoceros populations, Okita-Ouma *et al*. (2010) followed the procedure of Cromsigt *et al*. (2002) and found that only the exponential model returned sufficiently precise estimates of model parameters. The time frame for both studies was roughly ten years. Chamaillé-Jammes *et al*. (2008) used AIC_c_ to compare several discrete-time scalar models of population dynamics with multiplicative error, including the genRicker, for aerial survey counts of an elephant (*Loxodonta africana*) population exhibiting several years of considerable growth after cessation of culling, followed by fluctuations, and obtained the genRicker as best fit with *θ* = 6.55 (SE = 2.51) but ultimately found that only a model with *K* related to rainfall adequately explained their data. We propose that an extended period of growth indistinguishable from exponential growth may be common for expanding populations of megaherbivores. In addition to the SKKR population and those of Okita-Ouma *et al*. (2010), Knight *et al*. (2001) and Gough and Kerley (2006) reported exponential growth for expanding populations of black rhinoceros and elephant, respectively, while Brodie *et al*. (2011) deduced density independent vital rates from mark-recapture survey data for a black rhinoceros population recovering from poaching.

Our estimate of the (annual) intrinsic rate of growth for the SKKR population of 0.102 ± 0.017 is at the high end of the range of published values with estimates typically below 0.1 (Owen-Smith, 1988; Knight *et al*. 2001; Okita-Ouma *et al*. 2010; Brodie *et al*. 2011; Ferreira *et al*. 2011; Greaver *et al*. 2014); Cromsigt *et al*. (2001) obtained *r* = 0.1 while Okita-Ouma *et al*.’s (2010) largest estimate was 0.086 ± 0.022. The scaling law *r* = 1.5 *W*^0.36^ (Caughley and Krebs 1983), where *W* is mean adult live weight in kilograms, yields *W* = 1750 for *r* = 0.102 and higher rates for lower *W*, so the estimate of SKKR is within theoretical expectations as *W* lies in the range 700 – 1400 for black rhinoceros (Owen-Smith, 1988; 1000kg yields *r* = 0.125).

Since the population grew exponentially, we built a stage-based matrix model employing biological states of calf, subadult, and adult (Law and Linklater 2014; Table 1). As the final planned introduction occurred in Dec-97, we employed this matrix model to examine the structured population dynamics from Dec-98 through Dec-08, which period included the removal of five rhinoceros in May-06. The matrix-model projection of the SKKR population vector for Dec-98 (5 female calves, 11 female subadults, 8 female adults, 3 male calves, 11 male subadults, 4 male adults, Keyfitz’s Δ = 0.227 from the stable-stage distribution) converged monotonically and initially quite rapidly on the matrix model’s stable-stage distribution. In this sense, the matrix model and initial population vector for Dec-98 exhibited only mild transient dynamics. Moreover, the projected annual growth rates for Dec-98 through Dec-07 ranged from 1.0967 to 1.1267 (Table 9) versus the asymptotic rate of 1.1078. Also, the pattern of rankings for sensitivities and elasticities of the model’s asymptotic growth rate and of annual growth rates were similar; in particular, adult survival was the most influential parameter on both (Table 10). Thus, once reintroductions ceased, the deterministic dynamics implied that the SKKR population should approach its stable-stage-distribution dynamics within the lifespan of black rhinoceros. If these results are typical for black rhinoceros, then transient dynamics will largely reflect unusual population structure rather than the sub-dominant eigenvalues. Nevertheless, the computation of projected annual growth rates from a matrix model could provide a useful tool for managers planning a removal of individuals to check for a robust population structure and avoid unintended low short-term growth rate in response to the removal, viz., values below the asymptotic rate indicate a less robust population structure.

On the other hand, the actual SKKR population vectors, Dec-99 through Dec-08, did differ from the matrix model projections and did not converge monotonically on the model’s stable-stage distribution, exhibiting instead small fluctuations (Fig. 5). Similarly, Keyfitz’s Δ between the annual projections of the SKKR population vectors for each December, 1998– 2007, and the SKKR population vectors for the following December, though typically less than 0.1 did not converge on zero over this period (Fig. 6). Thus, the deterministic model did not capture the structured population trajectory in all detail. As noted above, we previously found no influence of rainfall on SKKR demography and here estimated environmental stochasticity as absent. Direct examination of the structured population trajectories and matrix projections (Table 8) indicated that the discrepancies could largely be attributed to the deviations between the actual numbers of, and model projections of, sex-specific recruitment each year. We previously found no deterministic explanation for the variation in interbirth intervals or birth sex in the SKKR population (Law *et al*. 2013, 2014) in terms of biologically plausible covariates and proposed that the variation was due to demographic stochasticity.

As measured by *R*^2^ (Table 3), the fit of the scalar models to the abundance data was extremely high, yet the better fit of the discrete-time exponential model compared to the continuous-time model indicated the presence of process noise in the dynamics. From the unstructured population, we estimated environmental stochasticity to be negligible. The climate of the study area is warm temperate (Fike 2011) with rainfall expected to be the main driver of environmental influence on dynamics. Yet we found no evidence for influence of rainfall on interbirth intervals, age at first reproduction, or birth sex (Law *et al*. 2013, 2014), consistent with the estimation of no environmental stochasticity. In combination, this evidence of process noise manifest as variation in fecundity and birth sex (there was very little mortality, Law *et al*. 2013) in the SKKR population dynamics in the absence of environmental influence supports our interpretation of this variation as due to demographic stochasticity.

There have been few estimates of demographic stochasticity for long-lived vertebrates, or even of mammals. Using unstructured population models, our estimate 0.127 (or 0.178 for the female segment only) of demographic stochasticity for SKKR compares with values of 0.267 for a population of Swiss ibex (*Capra ibex*) (Sæther *et al*. 2007b), 0.28 for the Soay sheep (*Ovis aries*) of Hirta Island, U.K. (Lande *et al*. 2003, Table 1.2), 0.571 for Scandinavian wolverines (*Gulo gulo*) (Sæther *et al*. 2005), and 0.745 for a population of Norwegian roe deer (*Capreolus capreolus*) (Grøtan *et al*. 2005), the last being the highest value for demographic stochasticity reported for a mammalian population by any method. For structured population models, our estimate of 0.204 compares with 0.084 for a population of wandering albatross (*Diomedea exulans*) (Engen et al. 2005), 0.15 for a Norwegian island population of moose (*Alces acles*) using the method of Engen *et al*. 2005 (Sæther *et al*. 2007a), 0.155 and 0.180 for two populations of Scandinavian brown bear (*Ursus arctos*) (Sæther *et al*. 1998b). The studies of Sæther *et al*. (1998b), (2007a) and (2007b) also reported very low (< 0.008) to negligible values for environmental stochasticity *σ*_*e*_^2^.

Our estimates of demographic stochasticity are similar to those for species with long generation times (Sæther *et al*. 2007a); the larger values obtained in the studies cited above appear to result from a combination of high adult survival and variable recruitment. For the SKKR population, demographic stochasticity manifested itself in random variation in IBIs (Law et al. 2013) and also birth sex (Law et al. 2014) with the difference between actual population structure and matrix-model projections due differences between actual, and projected, sex-specific recruitment. Thus, the SKKR population is a more modest example of high adult survival and variable recruitment. The absence of environmental stochasticity and moderate demographic stochasticity implies that the mean of observed ln(*N*_*t*+1_/*N*_*t*_) values should not differ too much from the intrinsic rate of growth, which no doubt aided the success of this reintroduction. Calf mortality was essentially absent for SKKR, but important in the studies of Hrabar and du Toit (2005), Brodie et al. (2011), and Greaver et al. (2014) and may be contributions to environmental and/or demographic stochasticity and/or the deterministic dynamics of those populations.

The form of density dependence for megaherbivores remains uncertain, though an extended period of exponential growth may be common. Whether this period of growth continues until a threshold followed by a sharp decline in pgr, or to an inflexion followed by a more gradual reduction, requires further study with time series of census data over the full range of population sizes. We previously reported an increase of age at first reproduction in SKKR with increasing population size (Law et al. 2013) despite no apparent resource limitation (van Lieverloo et al., 2009) and suggested this response was socially mediated (Bronson 1989:163). Increase in age at first reproduction might therefore provide a practical early warning sign of density feedback in an expanding population of megaherbivores prior to detectable slowing in growth rate. Further study across populations and species is required to explore this possibility. Demographic contributions to process noise are likely important for understanding megaherbivore population dynamics, as such populations are often relatively small, and therefore of relevance to both the impact of removals on donor populations and on the performance of reintroduced populations. Though megaherbivores may exhibit some robustness to environmental variation, we would not expect the absence of environmental stochasticity observed in the SKKR dynamics to be typical of megaherbivores, especially given the diverse habitats occupied by black rhinoceros and elephant in particular. Hrabar and du Toit (2005), Gough and Kerley (2006), Chamaillé-Jammes *et al*. (2008) and Lee *et al*. (2011) all reported influences of rainfall on the demography of megaherbivores, black rhinoceros in the first instance and African elephant in the other three. Nevertheless, Brodie *et al*. (2011:355) found no temporal variation in vital rates, over a 14-year period, of the black rhinoceros population they studied in the ‘most extreme desert-dwelling ecotype of black rhino’. It would be interesting to know the rainfall pattern over that time.

Given adequate monitoring of a potential donor population to avoid unreliability due to estimates of abundance rather than census data (Ludwig 1999), one might employ the model of Dennis *et al*. (1991) to compute probabilities of extinction (or of an unacceptable reduction in numbers) prior to the inception of managed removals as a check on the robustness of the population. Engen *et al*. (1998), however, pointed out that the model of Dennis *et al*. (1991) neglects demographic stochasticity, which could result in biased estimates of such calculations. Knight *et al*. (2001) obtained from the model of Dennis *et al*. (1991) the probability of a particular black rhino population decreasing from only 33 to ten individuals to be negligible. Similar calculations using both the model of Dennis *et al*. (1991) and that of Engen *et al*. (2005), which incorporates demographic stochasticity, also yielded negligible probabilities for various scenarios with the SKKR population and the differences in results between the two methods were of no practical consequence. When such probabilities are not negligible, the differences in the models may not be unimportant, e.g., for populations influenced by both environmental and demographic stochasticity, especially if the latter also influences survival, and not just fecundity as for SKKR. In formula (45), derived from Engen *et al*. (2005), the contribution from a given nonzero entry of the projection matrix depends on the dominant eigenvalue, the stable-stage distribution, the sensitivity of *λ* to that entry, and the variance of the demographic parameter the entry is the mean of. The importance of demographic stochasticity is also evident in the form of founder effects of course, so selection of individuals for removal is important for both the donor and target populations.

## Acknowledgments

Our collaboration is a by-product of an International Science Liaison Foreign Fellowship, National Research Foundation, Republic of South Africa, PRL shared with Wayne Linklater (WL) in 2005. We thank: WL, for his pivotal role in obtaining this fellowship, bringing the 3 authors together, and for ongoing dialogue on rhinos; Graham Kerley, for hosting WL and PRL during their fellowship at the Centre for African Conservation Ecology, Nelson Mandela Metropolitan University; and the SKKR field rangers.

